# Synaptic networks shape clinical phenotypes in neurodevelopmental disorders: An integrative clinical, genetic and biological perspective

**DOI:** 10.64898/2026.01.23.701319

**Authors:** Juliana Ribeiro-Constante, Julia Romagosa-Perez, Alfonso Oyarzábal, Enric X. Martin Rull, Angeles García-Cazorla

**Affiliations:** Neurometabolic Unit and Synaptic Metabolism Laboratory, Neurology Department, CIBERER and MetabERN, Sant Joan de Déu (SJD) Children’s Hospital, Barcelona 08950, Spain; Institut de Recerca Sant Joan de Déu, Barcelona 08950, Spain; Pediatric Neurometabolism and Personalized Therapies Lab, University Abat Oliba CEU, Barcelona 08022, Spain; Automatic Control Department, Technical University of Catalonia (UPC), Campus Diagonal Sud, Barcelona 08028, Spain; Medicina i Recerca Translacional, Facultat de Medicina i Ciències de la Salut, Universitat de Barcelona, Barcelona, 08036, Spain

**Author notes:** Corresponding author: Ángeles García-Cazorla, Corresponding author’s address: Neurometabolic Unit and Synaptic Metabolism Lab, Department of Neurology, Hospital Sant Joan de Déu and Institut de Recerca Sant Joan de Déu, CIBERER-ISCIII and MetabERN, Corresponding author’s, Corresponding author’s. These authors contributed equally to this work (Juliana Ribeiro-Constante, Julia Romagosa-Perez).

**Keywords:** Human Phenotype Ontology, synaptic biological processes, developmental gene expression, synaptic compartment localization, neurodevelopmental disorder clustering

## Abstract

**Background:** Clinical variability in synaptopathies suggests complex multidimensional interactions. The resulting phenotypes depend not only on the affected gene, but also on its functional role, subcellular localization, and spatiotemporal brain expression, among other potential influencing factors. Here, we present a proof of concept integrating genetic, synaptic, and developmental expression data to explore how shared pathways can lead to similar clinical manifestations across different synaptic disorders.We systematically curated published clinical data from 1,648 patients with pathogenic variants in 20 synaptic genes, through deep phenotyping using standardized Human Phenotype Ontology (HPO) annotation. Functional gene groups were defined based on synaptic localization, biological process, brain expression patterns and disease onset. Brain developmental expression data were obtained from the BrainSpan Atlas, normalized, and analyzed to assess gene co-expression and timing of peak expression.

**Results:** Integrating synaptic biology with developmental gene expression data reveals phenotypic emergence through interconnected synaptic characteristics. Subcellular localization shapes manifestations: presynaptic variants associate with motor disorders and postsynaptic variants with broader systemic involvement. Biological processes further refine phenotypes, with transport processes associated with hypotonia and signaling pathways linked to early-onset symptoms. Developmental timing adds dimension: prenatal expression peaks align with cognitive impairments and postpubertal peaks with motor decline. Crucially, clinical profiles mirror co-expression patterns of affected genes, demonstrating dynamic interactions across spatial, functional, and temporal dimensions.

**Conclusions:** By mapping symptoms into synaptic networks across developmental windows, this work establishes the foundation for a “SynGO-Clinic” platform, a clinical extension of SynGO, to advance biological insight, diagnostics, and targeted therapies in neurodevelopmental synaptopathies.

## Background

Neurodevelopmental disorders exhibit a high degree of clinical heterogeneity; however, a growing number of studies have shown that many of these diseases share a common origin in synaptic dysfunction. The synapse, a fundamental structure for neuronal communication, relies on highly specialized molecular machinery. Genetic alterations affecting this machinery can lead to a broad spectrum of neurological symptoms, whose profiles depend not only on the gene involved but also on multiple interrelated factors.^1^

The subcellular localization of the protein (presynaptic/postsynaptic),^2,3^ its role in processes such as vesicular trafficking^4^, its regional brain expression,^5,6^ and its temporal activity across developmental stages^7,8^ collectively shape the clinical phenotype. Consequently, mutations in distinct genes may lead to similar symptoms when they disrupt shared synaptic protein networks.^9^

In recent years, it has become evident that the traditional gene-centric approach is insufficient to fully understand neurodevelopmental disorders. Instead, an integrative framework has emerged, emphasizing gene networks, spatiotemporal dynamics, and environmental interactions.^10,11^ Within this paradigm, distinct neurological conditions may converge through shared functional modules in common regulatory networks.^12^

Inspired by the work developed in the SynGo platform,^13^ we aimed to include clinical aspects in this integrative perspective. Therefore, this article presents a proof of concept based on the integration of genetic, synaptic, and developmental expression timing data to explore how shared pathways can lead to similar clinical manifestations across different synaptic disorders. The central hypothesis is that the phenotype does not depend solely on the affected gene, but on the broader biological context in which that gene operates. This work is in fact a pilot study for a future fully developed “SynGo-Clinic platform” that could be useful for accurate clinical definitions and diagnosis, their connection with biological pathways and targeted therapies. This platform is aimed to be a dynamic intelligent tool nourished by continuous novel information derived from newly published patients.

## Results

### Curated clinical dataset derived from systematic literature review

A comprehensive literature review was conducted, covering more than 307 publications, including 196 case reports (less than 5 patients reported) and 111 case series, published between 1994 and 2024. Although the primary search strategy focused on studies from the past five years using PubMed, older publications were included when they described previously reported patients relevant to the analysis. For each gene, an average of 220 papers were identified (16 to 771 papers per gene). The goal of this review was to extract the most detailed clinical and genetic information available for each reported patient carrying a pathogenic variant in one of the analyzed genes. A summary of the review process is provided in Supplementary Table 1.

Once the review was completed, the dataset included 1,862 patients and more than 17,900 HPO terms. To ensure data homogeneity, a maximum threshold of 100 patients per gene was applied. After filtering, the final dataset comprised 1,648 patients and more than 1,281 unique HPO terms (Table 1).

**Table 1.** Demographic table of cohort characteristics. *(Separate file)*

The extracted data included: a) The specific genetic variant reported per patient; b) Age at onset of the first symptoms (when available); c) Age at assessment (when available), d) Ethnicity (when reported); e) HPO-coded symptoms derived from the clinical descriptions provided in each article.

Reference to the article where each patient was reported was also recorded. This curated dataset forms the basis for the analyses presented in subsequent sections. At this stage, the complete dataset is not publicly available but is accessible upon reasonable request from the corresponding author.

### Synaptic genes linked to neurological disorders are associated with a highly diverse range of symptoms

Table 1 presents the demographic and clinical characteristics of the cohort, comprising 1,648 patients with pathogenic variants in 20 synaptic genes. Each genetic group included a mean of 80 patients, with a minimum of 58 and a maximum of 101 (Supplementary Figure 1A). The mean age at clinical description was 14.51 years (range: 0–47), and 54% of patients were male. However, sex distribution varied substantially. CDKL5 and CASK showed strong female predominance (female ratios of 66% and 70%, respectively), likely reflecting their X-linked inheritance patterns. Conversely, FMR1 and OPHN1 had a male predominance of over 85%, and SLC6A8 reached 80% male, in line with known X-linked recessive traits (Supplementary Figure 1 (B and D)).

On average, patients presented with 10 distinct HPO terms (range: 3.7–24). A total of 1,281 unique HPO terms were collected across the cohort, underscoring the clinical heterogeneity associated with synaptic gene dysfunction.

Genes such as CASK, CDKL5, PURA and SYNGAP1 showed the highest phenotypic burden, with mean HPO terms per patient of 13.76, 13.20, 13.33 and 18.25, respectively. In contrast, mutations in genes like LGI1, and DNAJC5 were presented in older individuals (mean ages 42.52, and 44.79 years) (Supplementary Figure 1C) and later onset (means of 7.96 and 5.17), and were associated with fewer symptoms per patient.

Each gene was categorized according to its biological role and synaptic localization using the SynGO ontology. These classifications are visualized in Figure 1 and detailed in Table 1. This classification establishes the foundation for subsequent analyses linking molecular function and synaptic location to specific phenotypic patterns. No significant differences were observed in the number of HPO terms per patient between functional SynGO categories (*p* = 0.38). To reduce heterogeneity, we applied an aggregation strategy based on HPO parent terms. A customized aggregation scheme was used (Supplementary Table 2), which prioritized specificity for neurological symptoms while generalizing non-neurological features. This approach followed the HPO tree structure to ensure ontological consistency. After aggregation, we quantified intragenic phenotypic similarity before and after term grouping. As shown in Supplementary Figure 2 (A and B), the use of aggregated terms resulted in increased intragenic similarity across most genes.

**Figure 1.**
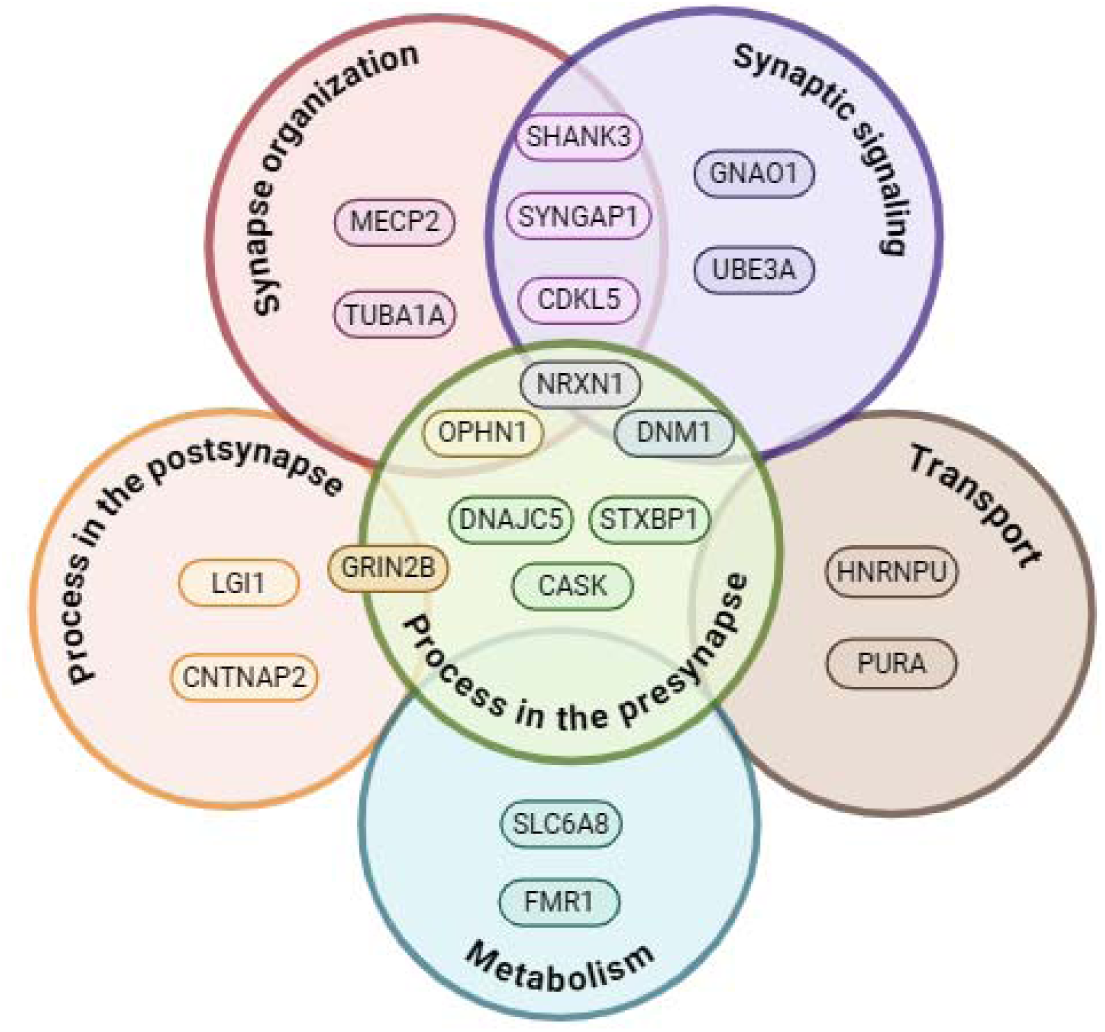
Genes included in the study grouped by SynGO-defined biological processes. This figure presents the set of synaptic genes analyzed in the study, classified according to their associated biological processes as defined by the SynGO ontology. The selected genes represent a broad range of functions involved in synapse and diverse neurodevelopmental and neurological disorders.

In Supplementary Figure 3, the twenty most frequent symptoms across the entire cohort are shown. As expected, symptoms such as epilepsy, abnormal communication, intellectual disability, and neurodevelopmental delay consistently define the general phenotypic profile associated with synaptopathies. However, among the most frequent features, some non-neurological symptoms were also identified, including abnormalities of the head or neck, musculoskeletal abnormalities, and digestive system disorders. In total, standardized HPO annotation captured 1,281 unique terms, highlighting the remarkable clinical diversity arising from synaptic gene mutations.

### SynGO-defined biological processes and synaptic localization influence symptom presentation

We systematically compared symptom profiles across genes stratified by SynGO-defined biological processes and synaptic localization (gene classification shown in Supplementary Figure 4). Figure 2 illustrates the 15 most prevalent symptoms within each biological process group, revealing distinct phenotypic patterns. As an example, Supplementary Figure 5 shows some symptoms with statistically significant variation across groups (p < 0.05, Fisher’s exact test).

**Figure 2.**
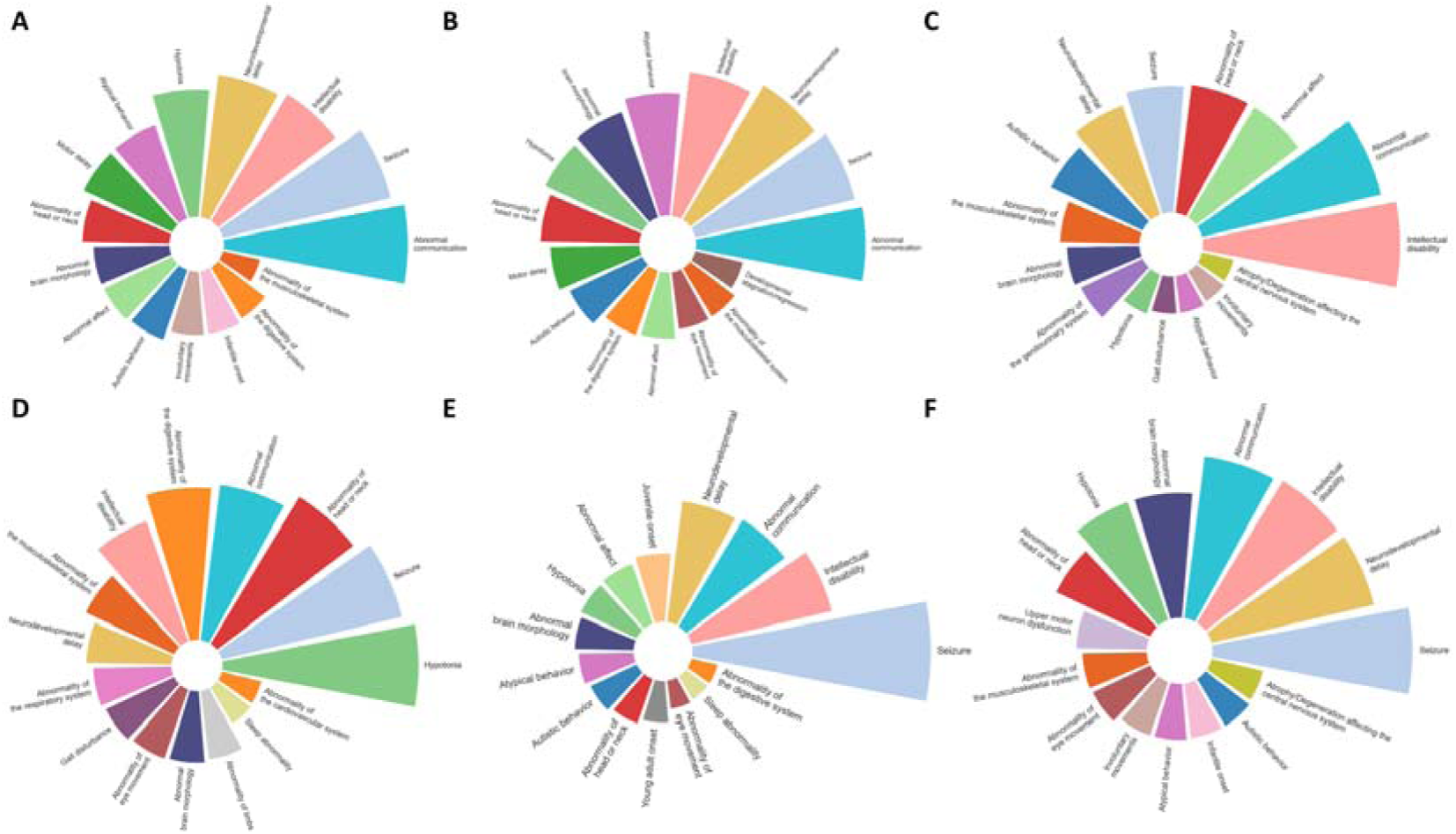
Distribution of the 15 most frequent symptoms within each biological process group. Symptoms were identified based on their relative frequency within each group. A) Synaptic signaling (n = 601 patients); B) Synapse organization (n = 633 patients); C) Metabolism (n = 141 patients); D) Transport (n = 181 patients); E) Processes in the postsynapse (n = 217 patients); F) Processes in the presynapse (n = 536 patients).

Following the criteria outlined in section “Symptom Frequency Analysis”, symptoms were assigned to a biological process. Table 2 summarizes the characteristic symptoms significantly associated with each biological process. The Transport group showed the highest number of enriched symptoms, including *hypotonia* (78.45%, *p* < 0.001), *abnormality of the head or neck* (68.5%, *p* < 0.001), and *musculoskeletal abnormalities* (39.22%, *p* < 0.001). This group also showed associations with non-neurological domains, such as *cardiovascular abnormalities* (16.57%, *p* < 0.001) and *integumentary system abnormalities* (14.9%, *p* < 0.001), highlighting a broader systemic involvement.

**Table 2.** Symptoms significantly enriched in each biological process group defined by SynGO annotation.

| Biologic process | Symptom | Frequency (%) | P-value |
| --- | --- | --- | --- |
| Synapse organization | Recurrent maladaptive behavior | 34.3 | < 0.001 |
|  | Developmental stagnation/regression | 17.69 | < 0.001 |
|  | Motor delay | 32.1 | < 0.001 |
| Synaptic signaling | Infantile onset | 23.62 | < 0.001 |
|  | Involuntary movements | 23.79 | < 0.001 |
|  | Motor delay | 36.77 | < 0.001 |
| Transport | Abnormal myelination | 12.7 | < 0.001 |
|  | Abnormality of the musculoskeletal system | 39.22 | < 0.001 |
|  | Abnormality of head or neck | 68.5 | < 0.001 |
|  | Abnormality of the cardiovascular system | 16.57 | < 0.001 |
|  | Abnormality of the integument | 14.9 | < 0.001 |
|  | Abnormality of limbs | 27.6 | < 0.001 |
|  | Childhood onset | 16.57 | < 0.001 |
|  | Hypotonia | 78.45 | < 0.001 |
|  | Growth abnormality | 16.57 | 0.002 |
| Process in the presynapse | Upper motor neuron dysfunction | 22.01 | < 0.001 |
|  | Hypertonia | 6.71 | < 0.001 |
|  | Neonatal onset | 10.63 | < 0.001 |

Genes were grouped according to their involvement in specific synaptic biological processes. For each group, symptoms significantly more frequent compared to other groups were identified using chi-squared or Fisher’s exact tests, with Bonferroni correction applied for multiple comparisons. The table reports the enriched symptoms, their relative frequency (%) within the group, and the corresponding adjusted *p*-values. Only symptoms meeting the significance threshold (*adjusted P* < 0.05) and passing the group-level dominance criteria (see Methods, Section “Symptom Frequency Analysis”) are shown.

The Synaptic signaling group was characterized by *motor delay* (36.77%, *p* < 0.001), *infantile onset* (23.62%, *p* < 0.001), and *involuntary movements* (23.79%, *p* < 0.001), indicating early and dynamic motor involvement.

Similarly, the Synapse organization group was associated with *recurrent maladaptive behavior* (34.3%, *p* < 0.001), *developmental stagnation or regression* (17.69%, *p* < 0.001), and *motor delay* (32.1%, *p* < 0.001).

Genes involved in presynaptic processes showed increased frequencies of *upper motor neuron dysfunction* (22.01%, *p* < 0.001), *hypertonia* (6.71%, *p* < 0.001), and *neonatal onset* (10.63%, *p* < 0.001).

Synaptic location was also found to influence symptom presentation. For this analysis, we kept only the genes which were expressed either only in the presynaptic space or only in the postsynaptic space. Genes expressed in both locations were excluded, as they did not allow for a clear comparison between groups. Similarly, genes expressed in other synaptic locations were not considered, since each of those categories included only a single gene in our dataset, preventing meaningful group-level comparisons. Therefore, genes in the presynaptic space group are: DNAJC5, DNM1, CNTNAP2, NRXN1, OPHN1 and STXBP1, whereas genes in the postsynaptic group are: SYNGAP1, PURA, SHANK3, CDKL5, SLC6A8. The two presynaptic and postsynaptic groups were balanced in size, with 444 patients carrying mutations in presynaptic genes and 476 in postsynaptic genes. Figure 3A and 3B display the relative frequency of symptoms associated with presynaptic and postsynaptic genes, respectively. Symptoms that differed significantly between the two groups are indicated with an asterisk. Figure 3C highlights the specific symptoms that showed statistically significant differences between the two synaptic locations.

**Figure 3.**
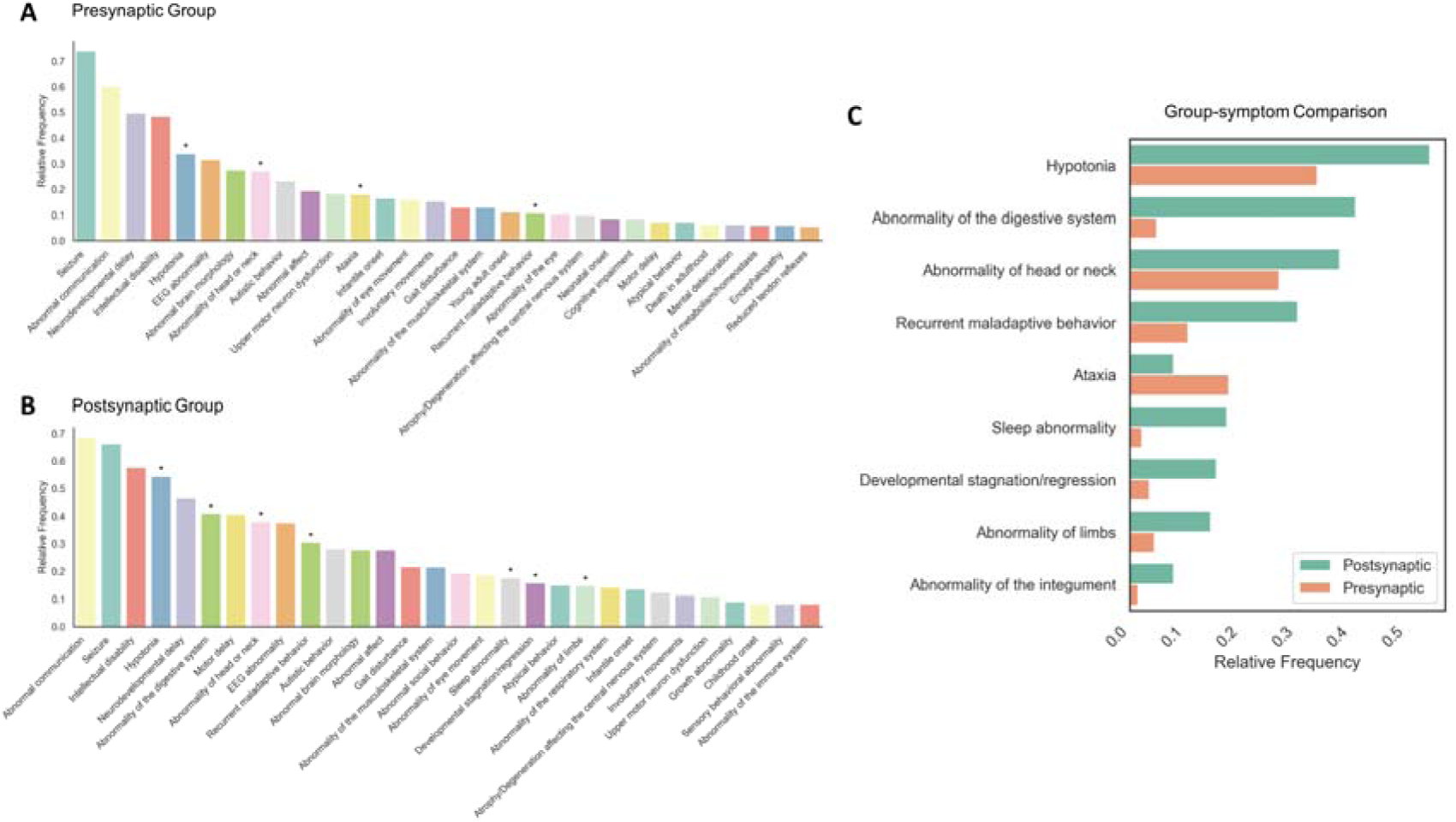
SynGO synaptic localization influences symptom presentation. **A**) Most frequent symptoms observed in patients with variants in presynaptic genes. B) Most frequent symptoms associated with postsynaptic gene variants. C) Symptoms showing significant differences in relative frequency between presynaptic and postsynaptic groups. Statistical differences in symptom prevalence between groups were assessed using chi-squared test or Fisher’s exact test, depending on expected cell counts. Bonferroni correction was applied to adjust for multiple comparisons. Symptoms with statistically significant differences are indicated by asterisks: p < 0.05 (*). Genes in the presynaptic space group are: DNAJC5, DNM1, CNTNAP2, NRXN1, OPHN1 and STXBP1. Genes in the postsynaptic group are: SYNGAP1, PURA, SHANK3, CDKL5, SLC6A8.

Mutations in presynaptic genes were primarily associated with *ataxia* (18%, *p* < 0.001), while mutations in postsynaptic genes were more frequently linked to *developmental stagnation or regression* (15.7%, *p* < 0.001), *hypotonia* (54.41%, *p* < 0.001), *digestive system abnormalities* (40.96%, *p* < 0.001), and *sleep abnormalities* (17.64%, *p* < 0.001), among others.

### Developmental gene expression patterns shape common neurological symptoms

Information regarding brain genetic expression across specific regions was also retrieved from the BrainSpan project. Brain expression matrices were constructed for each gene and developmental timepoint (Supplementary Figure 6). Pearson biserial correlations were calculated between gene expression profiles at each developmental age to identify gene pairs with similar expression trajectories (Supplementary Table 3).

Notably, DNM1 and STXBP1, as well as CNTNAP2 and NRXN1, showed strong correlations across developmental stages. Specifically, DNM1 and STXBP1 showed robust correlations during infancy (0–2 years; r = 0.855, *p* < 0.001), early childhood (3–7 years; r = 0.934, *p* < 0.001), post-pubertal (12.5–18 years; r = 0.908, *p* < 0.001), and adulthood (>18 years; r = 0.934, *p* < 0.001). Similarly, CNTNAP2 and NRXN1 demonstrated significant correlations in infancy (r = 0.791, *p* < 0.001), early childhood (r = 0.818, *p* < 0.001), pre-pubertal (8–12.5 years; r = 0.858, *p* < 0.001), post-pubertal (r = 0.814, *p* < 0.001) and adulthood (r = 0.82, *p* < 0.001). To complement these correlation analyses, we examined normalized mean expression values across brain regions (Table 3; non-normalized values in Supplementary Table 4). Notably, DNM1 and STXBP1 showed their lowest expression levels in the striatum, hippocampus, and thalamus, while CNTNAP2 and NRXN1 were least expressed in the cerebellar cortex, hippocampus, and striatum.

**Table 3.** Normalized mean brain expression across developmental age groups for selected pairs of correlated genes.

| Brain region | Mean expression |  |  |  |
| --- | --- | --- | --- | --- |
|  | DNM1 and STXBP1 |  | CNTNAP2 and NRXN1 |  |
| Amygdala | 0.81 | 0.72 | 0.78 | 0.69 |
| Anterior Cingulate and Medial Prefrontal | 0.68 | 0.68 | 0.79 | 0.73 |
| Auditory Association | 1.00 | 0.90 | 0.72 | 0.95 |
| Cerebellar Cortex | 0.83 | 0.99 | 0.06 | 0.26 |
| Dorsolateral Prefrontal | 0.77 | 0.79 | 0.87 | 0.84 |
| Early Auditory | 0.80 | 0.75 | 0.71 | 0.77 |
| Hippocampus | 0.50 | 0.37 | 0.39 | 0.36 |
| Inferior Frontal | 0.84 | 0.79 | 0.89 | 0.75 |
| Inferior Parietal | 0.86 | 0.78 | 0.81 | 1.00 |
| Lateral Temporal | 1.00 | 0.83 | 0.81 | 0.91 |
| Motor | 0.75 | 0.68 | 0.80 | 0.70 |
| Orbital and Polar Frontal | 0.86 | 0.83 | 1.00 | 0.89 |
| Primary Visual | 0.89 | 1.00 | 0.68 | 0.82 |
| Somatosensory | 0.90 | 0.87 | 0.84 | 0.75 |
| Striatum | 0.00 | 0.07 | 0.08 | 0.00 |
| Thalamus | 0.31 | 0.42 | 0.80 | 0.62 |
| **Correlation** | **0.933, $p < 0.001$** |  | **0.878, $p < 0.001$** |  |

Expression values were obtained from the BrainSpan Atlas and normalized across both samples and genes, ranging from 0 (lowest relative expression across regions) to 1 (highest relative expression). The Pearson correlation of mean expression across regions was r = 0.933, *p* < 0.001 for *DNM1* and *STXBP1*, and r = 0.870, *p* < 0.001 for *CNTNAP2* and *NRXN1*, indicating strong co-expression across brain development.

Building on these molecular and expression profiles, we next evaluated whether these patterns correlated with distinct clinical manifestations. As shown in Supplementary Figure 7, patients with DNM1 and STXBP1 variants predominantly presented motor abnormalities, including gait disturbances and involuntary movements. Conversely, CNTNAP2 and NRXN1 were more frequently associated with a neurodevelopmental symptom profile, including intellectual disability, developmental delay, impaired communication, and autistic behavior.

### Developmental timing of peak gene expression is associated with distinct symptom profiles

Brain gene expression data across developmental stages were obtained from the BrainSpan Atlas. For each gene, we analyzed expression trajectories across four developmental windows: prenatal, infant, childhood, and adulthood (Supplementary File 2). Genes were then grouped based on their age of maximum brain expression, resulting in three main categories expression groups: prenatal (CASK, SYNGAP1, OPHN1, HNRNPU, TUBA1A, UBE3A, GRIN2B, NRXN1), infant (SLC6A8, FMR1, LGI1, CNTNAP2, CDKL5, GNAO1) and post-pubertal and adulthood (STXBP1, DNAJC5, PURA, MECP2, DNM1) (Figure 4A).

**Figure 4.**
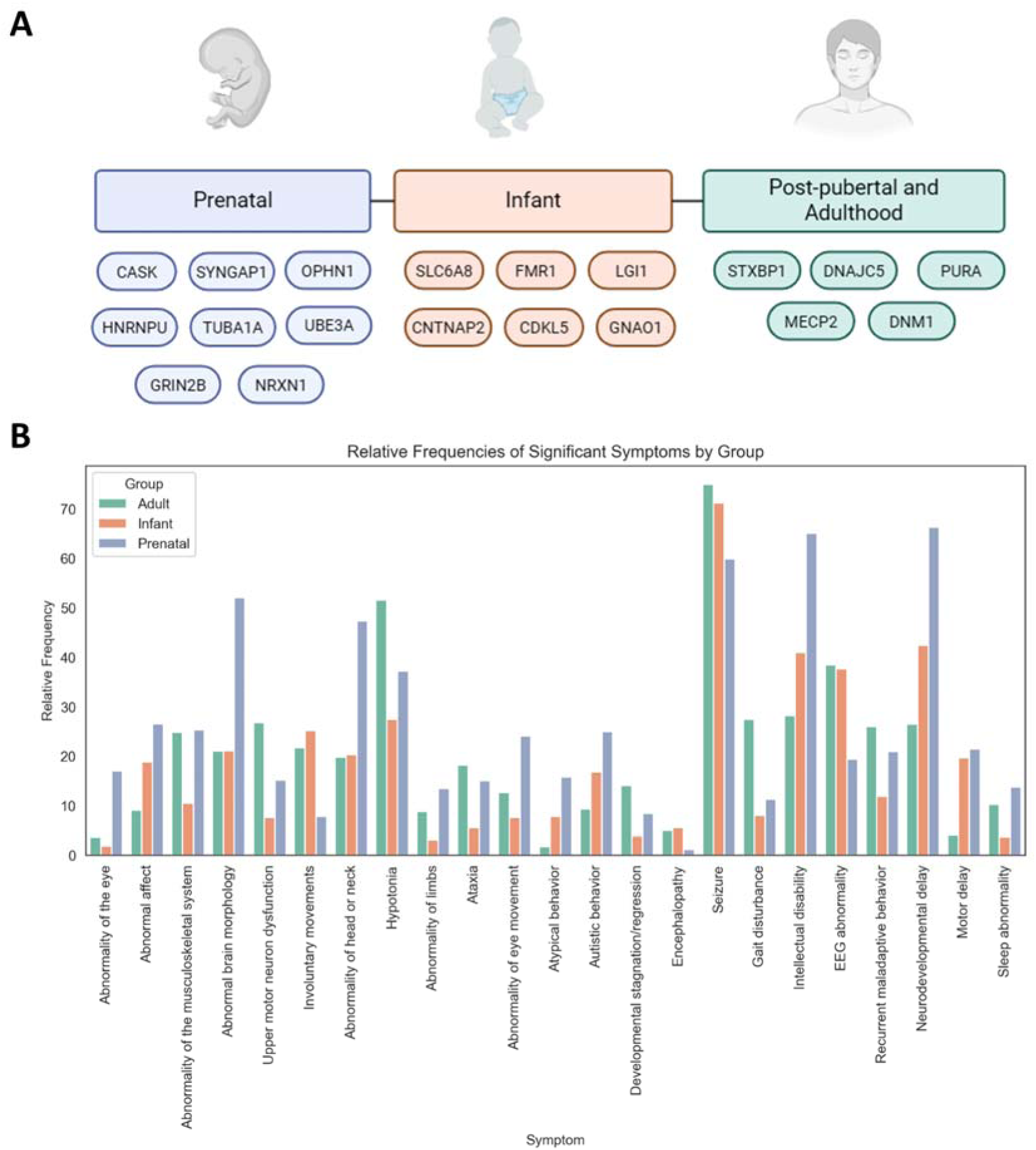
Characteristic symptoms associated with gene groups classified by their age of maximum expression across brain development. A) Schematic representation of gene groups based on their developmental peak of expression, as extracted from the BrainSpan Atlas. Developmental age groups were defined according to the GTEx resource (https://gtexportal.org/home/aboutdGTEx). B) Relative frequency of significant symptoms according to the gene’s peak expression stage (adult, infant, and prenatal). Only significantly different symptoms between groups are shown. Statistical significance was determined using Fisher’s exact test or chi-squared test. P-value was corrected using the Bonferroni method. Symptoms were associated with a group if the genes within that group collectively accounted for at least 30% of the total symptom frequency. Symptoms dominated by a single gene were excluded, following previously described dominance criteria.

It was then examined how symptom presentation varied across these developmental expression groups (Figure 4B). Overall, it is observed that moment of genetic brain expression also influences symptom presentation. Genes with prenatal peak expression were associated with a broader range of symptoms, including structural brain anomalies (52.03%, *p* < 0.001), hypotonia (37.18%, *p* < 0.001), neurodevelopmental delay (66.25%, *p* < 0.001), intellectual disability (65%, *p* < 0.001), and autistic behavior (25%, *p* < 0.001) (Supplementary Table 5). In contrast, genes with infant peak expression were more frequently linked to seizure-related phenotypes (71.16%, *p* < 0.001), encephalopathy (5.52%, *p* = 0.005), and behavioral abnormalities such as autistic behavior (16.77%, *p* < 0.001). Genes with adult peak expression showed higher association with motor symptoms, including gait disturbance (27.45%, *p* < 0.001), upper motor neuron dysfunction (26.73%, *p* < 0.001), involuntary movements (21.72%, *p* < 0.001), ataxia (18.14%, *p* < 0.001), and hypotonia (51.55%, *p* < 0.001).

Interestingly, although our dataset lacks precise age-of-onset data for individual symptoms, we inferred temporal patterns by analyzing symptom prevalence across age groups (see Methods). For genes with peak expression in adulthood (*STXBP1, DNAJC5, PURA, MECP2,* and *DNM1*), symptom profiles varied significantly with the patient’s age at evaluation (Supplementary Figure 8A and B). Specifically, *hypotonia* showed an inverse correlation with patient age, more frequently reported in infancy and early childhood, while *involuntary movements*, *musculoskeletal abnormalities*, *upper motor neuron dysfunction*, and *ataxia* became more prevalent in older age groups.

### Disease onset influences clinical profiles

Age at symptom onset was reported for a subset of patients (Figure 5). Most of the patients in our cohort report a symptom onset between 28 days and 1 year of age (218 patients), followed by onset between 1 and 5 years (117 patients), 15 to 40 years (93), 0 to 28 days (84) and 5 to 15 years (75). Only 6 patients presented symptom onset after the age of 40.

**Figure 5.**
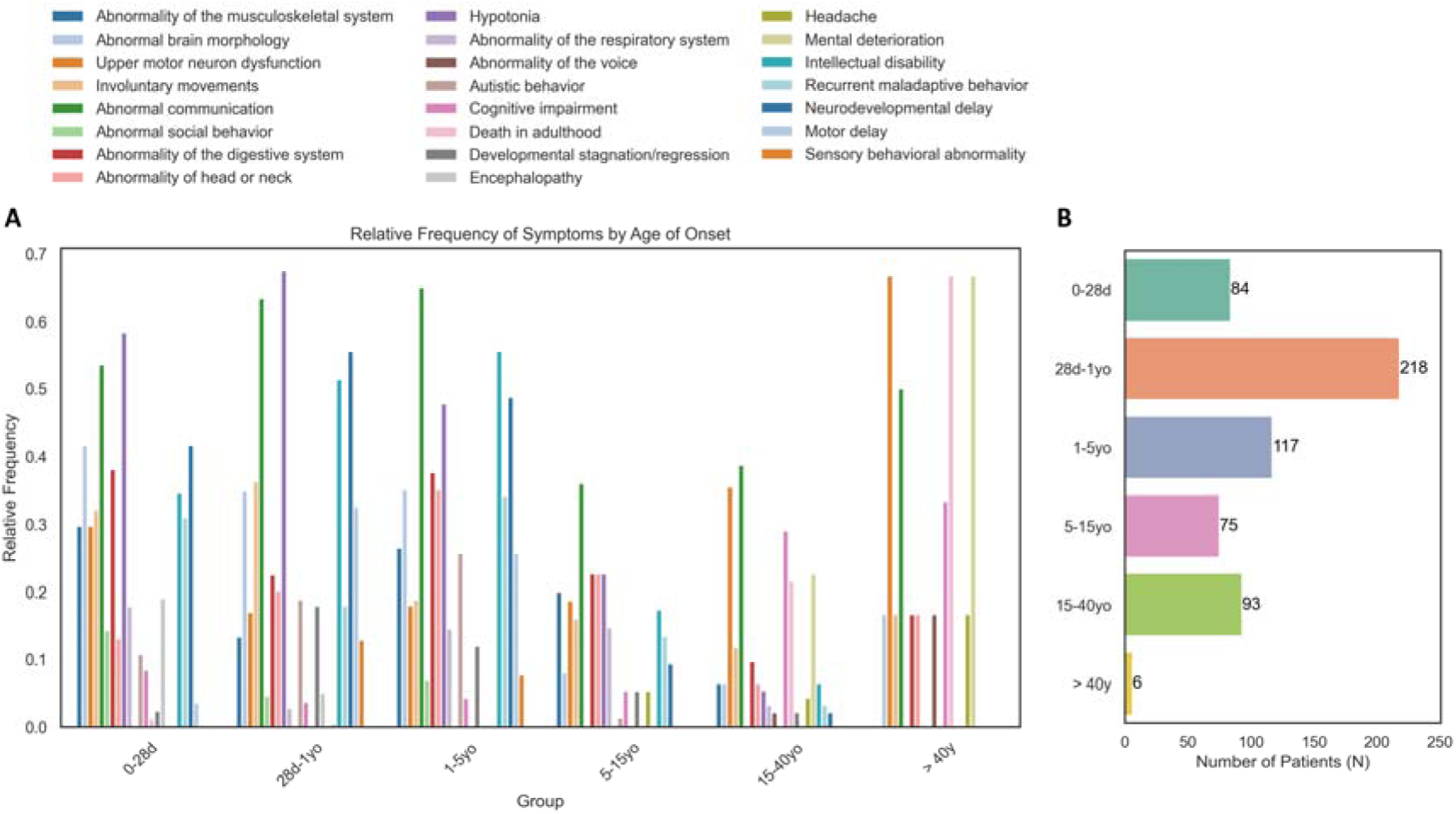
Symptoms with significantly different relative frequencies according to age at disease onset. **(A)** Displayed are symptoms whose prevalence varies significantly (Chi-square or Fisher test p-value<0.05) across onset of the diseases age groups (when reported): neonatal (0–28 days, N = 84), infant (28 days–1 year, N = 218), early childhood (1–5 years, N = 117), childhood (5–15 years, N = 75), young adulthood (15–40 years, N = 93), and late adulthood (>40 years, N = 6). Each bar represents the relative frequency of the symptom within the corresponding age group. Distinct clinical patterns emerge, such as higher frequencies of hypotonia, communication abnormalities, and neurodevelopmental delay in earlier onset groups, whereas symptoms like cognitive deterioration, sensory processing abnormalities, and adult mortality are more prevalent in older onset groups. **(B)** The right panel shows the number of patients in each age group, highlighting the underrepresentation of patients with disease onset after 40 years of age.

Patients with early-onset disease (before 5 years of age) more frequently exhibited hypotonia (0–28 days: 58.3%, 28 days–1 year: 67.4%, 1–5 years: 47.9%), abnormal communication (53.6%, 63.3%, and 64.9%, respectively), neurodevelopmental delay (41.7%, 55.5%, and 48.7%), and abnormal brain morphology (41.7%, 34.9%, and 35.0%).

In contrast, patients with late-onset disease (after 15 years) were more likely to show signs of mental deterioration (15–40 years: 22.6%, >40 years: 66.7%), cognitive impairment (29.0%, 33.3%), and upper motor neuron dysfunction (35.5%, 66.7%), including clinical features such as myoclonus, spasticity, and tetraplegia/paraplegia.

## Discussion

Despite the synapse being widely recognized as a critical hub in neurodevelopmental disorders,^18^ most prior studies have adopted a fragmented perspective, focusing on individual genes or isolated molecular pathways. This approach has limited our understanding of why mutations in distinct genes lead to overlapping clinical features, such as epilepsy or intellectual disability.^19^ Few studies have systematically examined how these genes interact with their synaptic, temporal, and functional contexts^10^ to produce convergent phenotypes, or divergent disorders due to developmental critical periods.^20.21^ Here, we integrate synaptic processes, localization, spatiotemporal expression patterns, and developmental trajectories to demonstrate that clinical phenotypes emerge from dynamic interactions within synaptic networks.

### Phenotypic diversity in synaptic genes

Neurodevelopmental disorders associated with synaptic genes exhibit phenotypic diversity extending beyond neurological symptoms, reflecting their roles in dynamic biological networks. These genes function differentially across development: for instance, SYNGAP1 regulates synaptic plasticity early in development^22^ and later modulates neuronal excitability,^23^ while UBE3A controls prenatal neuronal differentiation^24^ and postnatal synaptic homeostasis.^25^ As highlighted by Panagiotakos and Pasca,^21^ such spatiotemporal pleiotropy explains their multisystemic impact. This multifunctionality is evident in our cohort, which collectively spanned 1,281 unique HPO terms, including frequent non-neurological manifestations (musculoskeletal, digestive, and craniofacial anomalies) alongside core neurological features like intellectual disability and epilepsy (Supplementary Figure 3). These findings underscore that synaptopathies are systemic disorders, not merely localized synaptic dysfunctions.

### Synaptic processes and specific phenotypic profiles

Beyond subcellular localization, synaptic genes modulate clinical symptoms in a pathway-specific manner. When grouped by primary function—signaling, organization, transport, metabolism, presynaptic or postsynaptic processes—each category exhibited distinct phenotypic profiles (Figure 2, Table 2). These findings support the hypothesis that the nature of synaptic dysfunction critically shapes clinical outcomes, in line with previous conceptual models of pathway-specific circuit vulnerability.^26^

For example, synaptic organization genes were linked to maladaptive behaviors and motor delay, consistent with their role in maintaining synaptic architecture^3^ and network stability.^27^ In contrast, synaptic signaling genes were associated with involuntary movements and infantile-onset symptoms, reflecting disrupted excitation-inhibition balance. SHANK3 mutations impair synaptic stability,^27^ while UBE3A loss alters GABAergic transmission,^24^ collectively perturbing circuit maturation. Meanwhile presynaptic processes were predominantly associated with upper motor neuron (UMN) dysfunction and hypertonia in our analysis, suggesting that presynaptic alterations may contribute to motor dysfunction in neurodevelopmental disorders. In models of UMN hyperexcitability, synaptic remodeling, including presynaptic changes, has been observed as a downstream effect. Thus, presynaptic vulnerability could represent a convergent mechanism in disease progression.^28^ Notably, the group with the highest phenotypic specificity was synaptic transport genes (PURA, HNRNPU), which showed strong associations with hypotonia, abnormalities of head or neck, musculoskeletal defects, and limb abnormalities. These findings suggest convergent disruptions in synaptic trafficking^29^ shared pathways across high-demand tissues.^30^ PURA’s roles in mRNA transport and myelination,^31^ combined with HNRNPU’s regulation of synaptic genes via chromatin remodeling,^32^ may underlie this multi-system involvement.

### Influence of synaptic localization on clinical profile

The subcellular localization of synaptic genes significantly influences clinical manifestations. In our study, comparison of presynaptic (STXBP1, DNM1, DNAJC5, NRXN1, OPHN1, CNTNAP2) and postsynaptic genes (SYNGAP1, SHANK3, PURA, CDKL5, SLC6A8) revealed statistically significant phenotypic differences, highlighting compartment-specific effects (Figure 3). Consistent with synaptic biology principles^2^ and clinical reports (e.g., STXBP1^33^ and SYT1^34^), presynaptic gene mutations (e.g., STXBP1, DNM1) in our cohort predominantly associated with motor disorders, likely due to disrupted neurotransmitter release.^35,36^ In contrast, postsynaptic mutations exhibited broader phenotypes including neurodevelopmental (regression and hypotonia), systemic (digestive anomalies) and extracerebral (craniofacial malformations and integument abnormalities). This is consistent with literature emphasizing that postsynaptic dysfunction can affect not only synaptic transmission but also neurogenesis and cortical organization, as demonstrated for SYNGAP1, where non-synaptic pathological mechanisms have been identified as contributors to neurodevelopmental pathology.^37^ Furthermore, synaptic architecture and diversity, shaped by the molecular composition of postsynaptic complexes, are fundamental to the specificity of neuronal circuits and the emergence of complex behavioral phenotypes.^38,39^

### Developmental gene expression patterns shape common neurological symptoms

Gene expression patterns during brain development play a crucial role in the manifestation of neurological symptoms associated with synaptic genes. Our analysis, based on data from the BrainSpan project, revealed that the gene pairs DNM1–STXBP1 and CNTNAP2–NRXN1 exhibit sustained co-expression from neonatal stages into adulthood (*r* > 0.8, *p* < 0.05), with spatially similar patterns within each pair (Supplementary Table 4). However, both pairs show low expression levels in key brain regions (Table 3 and Supplementary Table 4).

Although DNM1 and STXBP1 exhibit low overall expression in the striatum, hippocampus, and thalamus (Table 3), regions critical for motor control,^40^ single-cell studies reveal their striking enrichment in GABAergic interneurons. These interneurons represent ∼15-20% of cortical neurons^41^ and ∼10-15% of striatal neuronal populations,^42^ where they modulate motor circuits through inhibitory signaling.^43^ In contrast, CNTNAP2 and NRXN1 are broadly low-expressed in the cerebellar cortex, hippocampus, and striatum (Table 3), regions linked to cognition and language, they display localized high expression in specific cell populations: Purkinje cells, which regulate motor coordination,^44^ and astrocytes, which modulate synaptic function in language-related circuits.^45.46^

These expression patterns are mirrored in clinical phenotypes. Patients with DNM1 or STXBP1 variants exhibit motor abnormalities (e.g., gait disorders, involuntary movements; Supplementary Figure 7A), whereas CNTNAP2 or NRXN1 mutations are linked to neurodevelopmental disorders (intellectual disability, autistic behaviors; Supplementary Figure 7B). These findings underscore how genes enriched in rare hub cell populations, such as GABAergic interneurons or astrocytes, can exert outsized effects on network function, where even limited dysfunction may breach compensatory thresholds, leading to pathology.^47,48^

### Developmental timing of peak gene expression is associated with distinct symptom profiles

The developmental stage of peak synaptic gene expression significantly influences symptom severity, onset, and progression. Our analysis revealed three distinct developmental trajectories (prenatal, infant, post pubertal adult), each associated with characteristic symptom profiles (Figure 4A and B). Notably, several genes with post pubertal-adult expression peaks are linked to early symptom onset,^49^ suggesting critical pre-peak functions through baseline activity or developmental roles. For example, STXBP1 mediates synaptic vesicle release at presynaptic terminals from prenatal stages, with mutations causing neonatal epileptic encephalopathies,^36^ while MECP2, despite adolescent peak expression, regulates gene networks essential for early brain development,^50^ and DNM1 governs synaptic vesicle endocytosis,^51^ with mutations leading to infantile epileptic encephalopathies.^52^ However, the relationship between gene expression and symptoms is not static; clinical manifestations often evolve as development progresses, reflecting either compensatory mechanisms or cumulative dysfunction in neural circuits.

Clinical symptoms evolve throughout development, with genes showing peak expression in post pubertal-adult stages often manifesting initially as hypotonia in early stages and progressing to complex manifestations (e.g., upper motor neuron dysfunction, ataxia) later in life (Supplementary Figure 8A and B). This evolution may reflect either: (1) A progressive degenerative process, as seen in DNM1- and PURA-related disorders where longitudinal studies document functional decline^29,51^ or (2) A maladaptive neurodevelopmental process where altered peak expression timing disrupts synaptic refinement during critical windows, leading to circuit disorganization that becomes apparent as network complexity increases.^20,21^ These processes may coexist in some disorders, with degeneration superimposed on earlier neurodevelopmental vulnerabilities.

### Symptom onset as a temporal axis in synaptic neurodevelopmental disorders

The age of symptom onset across multiple synaptic genes constitutes a fundamental dimension for understanding neurodevelopmental disorders. Our data reveal that specific clinical manifestations emerge preferentially at distinct developmental stages, reflecting dynamic interactions between synaptic dysfunction and nervous system maturation,^7,53,54^ While not strictly correlated with gene expression levels (Supplementary File 2), this temporal pattern shows clear functional associations when genetic alterations disrupt critical processes during vulnerable developmental windows.^55^

During the postnatal neonatal period (0-28 days), symptoms like hypotonia, seizures, and structural malformations dominate, indicating disturbances in neurogenesis and neuronal migration.^36,56^ From one month to five years, developmental delay, autistic behaviors, and cognitive impairments become prevalent, coinciding with cortical synaptic consolidation and pruning phases.^3,21^ From adolescence onward, late-onset manifestations including ataxia, motor deterioration, and upper motor neuron dysfunction emerge, frequently associated with genes maintaining adult expression (Figure 5A and B), suggesting roles in neuronal maintenance.^29,51^

This symptom chronology reveals core pathophysiological mechanisms: heightened vulnerability during critical periods,^20^ cumulative synaptic stress from persistent dysfunction,^57^ and progressive exhaustion of compensatory mechanisms.^58^ This framework explains clinical variability and could guide stage-specific interventions, from early plasticity modulation to late neuroprotection

## Conclusion

Our findings demonstrate that neurodevelopmental synaptopathies cannot be fully understood through isolated gene-level analyses but instead require an integrated framework encompassing synaptic localization, functional pathways, and developmental trajectories. Phenotypic convergence emerges from dynamic disruptions in shared biological networks, where spatial (subcellular compartments), molecular (functional modules), and temporal (critical windows) dimensions collectively shape clinical outcomes. This multidimensional perspective explains why mutations in distinct genes yield overlapping symptoms while also accounting for divergent phenotypes arising from developmental timing or circuit-specific vulnerabilities. By mapping these relationships, we provide a mechanistic roadmap for classifying synaptopathies, one that prioritizes biological context over isolated molecular signatures.

To bridge these insights into clinical practice, we propose a dedicated bioinformatics platform (Supplementary Figure 9) as a pilot “SynGo-Clinic” project, that stratifies patients based on synaptic processes, spatiotemporal gene expression, and phenotypic trajectories. Such a tool could enable mechanism-driven interventions, from early developmental plasticity modulation to late-stage neuroprotection, even in undiagnosed cases. Ultimately, this approach shifts the paradigm from gene-centric to network-centric understanding, offering a foundation for precision therapeutics in neurodevelopmental disorders.

### Limitations of the study

This study relies on aggregated clinical data from published sources, introducing heterogeneity due to variations in study methodologies and clinical assessments, which limits direct comparability and may affect phenotypic granularity. Publication bias is also possible, as severe or novel phenotypes are more frequently reported than milder or atypical cases. Furthermore, inconsistent documentation of symptom onset age constrains the ability to perform dynamic analyses of clinical progression in relation to gene expression patterns.

From an analytical perspective, co-expression analyses were limited to gene pairs with similar developmental profiles. Because the study has a cross-sectional design, it cannot track individuals over time or determine causal links between gene expression patterns and clinical phenotypes. Overcoming these limitations will require longitudinal clinical-genomic datasets, experimental validation in model systems, and causal inference methods.

Regarding resources and scope, functional gene categorization was based on SynGO, a synaptic-specific database that excludes non-synaptic roles (e.g., pleiotropic functions or extracerebral expression), which may obscure the interpretation of systemic or extracerebral phenotypes. Additionally, this pilot project focused on a limited number of genes, and no internal validation was performed due to its proof-of-concept nature. Future research should aim to replicate these findings in independent prospective cohorts and integrate large-scale clinical-genomic databases to enhance reproducibility and generalizability.

Collectively, these limitations highlight the need for prospective, standardized studies integrating genetic, phenotypic, and temporal data to validate and expand our findings.

## Materials and methods

### Creation of the database

#### SynGO ontology

Synaptic Gene Ontologies (SynGO)^13^ is a curated ontology based exclusively on experimental evidence from published studies, revised by experts. It includes 87 terms related to synaptic localization (cellular component) and 179 terms describing synaptic biological processes, encompassing a total of 1,602 genes.

SynGO distinguishes between synaptic location, referring to where a protein is active (cellular component), and biological process, describing the function it performs. At the highest level of the synaptic location hierarchy, synaptic proteins are categorized by their localization to the presynapse, postsynapse, synaptic cleft, extra-synaptic space, and synaptic membranes.

The relationships between terms reflect structural hierarchies, where smaller components are part of broader synaptic structures. Biological process terms follow the structure of the Gene Ontology (GO), with the main categories including pre- and postsynaptic processes, synaptic organization, synaptic signaling, axonal and dendritic transport, and metabolism as main terms with up to 6 levels of subclasses.

#### Inclusion criteria

To bridge the SynGO biological framework to the analysis of clinical phenotypes, we employed a selection criteria balancing biological coverage and clinical detail. First, we selected 2–7 genes from each of the six SynGO biological process categories (Presynaptic processes, Postsynaptic processes, Synaptic organization, Synaptic signaling, Metabolism and Transport; https://www.syngoportal.org/), ensuring that at least two genes were assigned a primary category to provide distinct functional coverage. Synaptic location was treated as an analytical grouping variable rather than an inclusion criterion.

Based on these data, we applied stringent clinical criteria to establish reliable correlations between phenotypes and biological processes. Each gene had to meet the following requirements:

- A well-documented association with human neurological disorders, as listed in OMIM (https://www.omim.org/).
- Published clinical data from at least 60 well-characterized patients.
- Each patient described with a minimum of three Human Phenotype Ontology (HPO) terms, ensuring phenotypic detail beyond the primary diagnosis.

To capture the breadth of synaptic pathophysiology, we included genes linked to both classical neurodevelopmental syndromes and other neurological conditions.

Genes satisfying all biological representation and clinical feasibility criteria formed the final set of 20 genes: MECP2, TUBA1A, STXBP1, SYNGAP1, CDKL5, CASK, DNM1, NRXN1, DNAJC5, OPHN1, UBE3A, SHANK3, GNAO1, SLC6A8, FMR1, GRIN2B, LGI1, CNTNAP2, PURA, and HNRNPU. Their classification by biological process and synaptic location is presented in Figure 1 and detailed in Supplementary Figure 4 and Table 1.

#### Data collection

A comprehensive literature review was conducted using the PubMed database. The search included the terms “Cohort of patients”, “Series of patients” and “Case report,” restricted to the last ten years (2014-2024). The aim was to identify studies that provide detailed clinical descriptions of patients with pathogenic or likely pathogenic mutations in any of the selected genes. We included all variant types, loss-of-function, gain-of-function, missense, deletions, and duplications, provided they were classified as disease-causing. This approach allows us to account for the possibility that distinct mutational mechanisms may converge on shared biological pathways, and to test the hypothesis that clinical phenotypes are shaped not only by the specific gene or variant involved, but by the broader biological context in which the gene operates.

The review process was conducted independently by two reviewers (JRP and JRC). The review methodology is detailed in Supplementary Table 1, which includes for each gene the list of included articles, number of patients per article, and the date of search. Studies were included if they reported detailed phenotypic and genetic data relevant to the predefined gene groups. The collected data were categorized into four main domains:

- Genetic Information: affected gene, variant details and corresponding protein
- Demographic Information: age, sex, and ethnicity (when available)
- Clinical Information: reported symptoms and age of symptom onset
- SynGO Information: assigned biological process and synaptic localization.

Once all the data were collected, we limited it to 100 maximum patients per gene and 60 minimum to maintain statistical coherence. When the number of patients exceeded 100 for a given gene, we selected the subset of individuals with the highest number of associated HPO terms, prioritizing richer phenotypic profiles. Additionally, patients described with fewer than three HPO terms were excluded from the analysis.

All clinical symptoms were annotated using the Human Phenotype Ontology (HPO) (https://hpo.jax.org). Specific HPO terms were assigned to each reported symptom to standardize terminology across studies, thereby enabling structured and reproducible analyses.

#### Symptom aggregation

To enhance the interpretability of phenotypic profiles, we aggregated detailed HPO terms into broader parent terms based on the HPO ontology hierarchy and the clinical experience of AGC and JRC. In order to preserve clinically relevant distinctions within the neurological domain, we defined a custom aggregation scheme that prioritizes specificity in neurological symptoms.

This refined grouping strategy was applied to homogenize HPO terms and reduce semantic redundancy, allowing for more consistent statistical comparisons across genes. The full mapping of original HPO terms to their corresponding aggregated categories is provided in Supplementary Table 2, including both the collected terms and their assigned parent groupings.

The methodology used to construct the database is outlined in Supplementary Figure 10, detailing the process followed from gene selection, literature review, HPO codification, data preprocessing, including enrichment of neurological symptoms, ending in the final dataset used to define the clinical hypothesis.

### Statistical analysis

#### Symptom Frequency Analysis

Phenotypic similarity between genes was assessed using the PyHPO Python library,^14^ which calculates similarity scores based on the HPO structure.

To identify differences in symptom prevalence between groups of genes, we applied either Chi-squared test or Fisher’s exact test, depending on the minimum expected cell count in the contingency tables. Statistical significance was defined as p < 0.05, and multiple comparisons were corrected using the Bonferroni method.

To investigate whether specific symptoms could be attributed to functional gene groups — defined by synaptic location, biological process, peak expression, and age of onset— we conducted a two-step analysis combining relative symptom frequency and gene dominance.

First, we aimed to identify symptoms that were significantly associated with each group. For this, we applied a proportional threshold based on relative frequency contributions. The relative frequency of each gene was normalized per symptom, and a symptom was assigned to a group if that group accounted for at least 30% of the total relative frequency of that symptom across all groups.

Next, to ensure that the observed associations were not driven by a single gene, we performed an additional dominance assessment. For each symptom, we calculated the relative frequency contributed by each gene within the group. A symptom was considered dominated if: (i) a single gene accounted for ≥70% of the total frequency, or (ii) in groups containing more than three genes, the top two genes together accounted for ≥80% (Supplementary Table 6). Symptoms meeting these criteria were annotated as dominated, and the contributing genes were recorded for downstream analyses.

This two-step approach allowed us to distinguish symptoms likely driven by the collective effect of multiple genes from those primarily influenced by individual gene effects, thereby reinforcing the validity of group-level phenotypic patterns.

#### Gene Brain Expression

Gene expression data were retrieved from the BrainSpan Atlas of the Developing Human Brain,^15^ covering multiple stages of brain development. The developmental transcriptome data^16^ contains data up to sixteen targeted cortical and subcortical structures across 42 brain specimens, spanning pre- and postnatal-development in both males and females.

##### Gene Co-expression Analysis Across Brain Regions and Development

Gene expression levels were analyzed in available cortical and subcortical brain structures in the BrainSpan Atlas. Gene expression levels were first normalized across samples and genes using Scaling with ranked subsampling method (SRS). Developmental age groups were defined according to the categories defined in GTEx resource^17^: infant (0-2 years), early childhood (3-7 years), pre-pubertal (8-12.5 years), post-pubertal (12.5-18 years) and adulthood (>18 years). For each developmental stage, expression matrices were constructed, in which columns represented genes and rows the mean expression across samples per region. This is illustrated as an example in Supplementary Figure 11A.

As gene expression in the brain depends on the developmental age, gene-co expression patterns were evaluated within each developmental group separately. Pearson correlation coefficients were computed between gene expression profiles within each developmental group. Gene pairs with a high correlation (r > 0.8, p < 0.05) across developmental stages were selected as co-expressed. As a result, pairs of genes which followed a similar expression in brain across different anatomical regions across development are obtained.

For each of these co-expressed gene pairs, we examined the clinical data to assess whether similar brain expression patterns correlated with overlapping symptom profiles. Specifically, it was compared the proportion of patients exhibiting each symptom within the gene pair group versus the rest of the cohort. Statistical significance was determined using chi-squared or Fisher’s exact tests, depending on data distribution, and multiple comparisons were corrected using the FDR method.

##### Time of maximum gene brain expression

Gene expression levels were first normalized across samples using SRS method. For each gene, the mean expression was computed at each age point using data from the BrainSpan Atlas. Locally weighted scatterplot smoothing (LOWESS) was used to characterize age-dependent expression trajectories. In addition, the spearman’s rank correlation coefficient (ρ) was calculated between gene expression and age. Based on these analyses, the age of peak expression was identified for each gene. Genes were then grouped according to their age of maximum expression into three developmental categories: prenatal, infant (0-3 years) and post-pubertal/adult (>18 years). None of the genes included in the study had peak brain expression during childhood. An example of the explained methods is described in Supplementary Figure 11B.

Therefore, for each group (prenatal, infant, post-pubertal/adult), we selected only those genes whose maximum expression fell within that developmental window. Statistically different symptoms between groups and associated symptoms were obtained following the same methodology described in methodology “Symptom Frequency Analysis”.

To test the association between symptom age and time of maximum gene expression, within each group, symptoms previously identified as significantly associated were tested for correlation with patient age. To assess the strength and direction of the relationship between binary symptom presence (0/1) and continuous age, we applied the point-biserial correlation coefficient. For each symptom, a correlation coefficient (r) and a p-value were calculated. Multiple testing corrections were applied using the Benjamini-Hochberg method (FDR).

Only symptoms with an adjusted p-value below 0.05 were considered significantly associated with age. These results were visualized using bar plots, showing both positive and negative correlations, thereby highlighting developmental patterns of symptom expression within gene groups characterized by similar expression timing.

## Declarations

### Ethics approval and consent to participate

Not applicable

### Consent for publication

Not applicable

## Availability of data and materials

The datasets generated is available in the [to be deposited] repository [DOI: to be deposited]. Any additional information required to reanalyze the data reported in this paper is available from the lead contact upon request.

## Author contributions

J.R.C., J.R.: conceptualization, data curation, formal analysis, methodology, investigation, interpretation of results, visualization, writing–original draft, writing–review and editing. A.O.: conceptualization, interpretation, writing–review and editing. A.G.C.: conceptualization, funding acquisition, supervision, interpretation, writing–review and editing. E.M.R.: conceptualization, supervision, methodology (bioinformatics platform), clinical framework design, writing–review and editing.

## Acknowledgements

We thank Matthjis Verhage for their valuable advice and input during the preparation of this work.

## SUPPLEMENTAL INFORMATION

**Supplementary File 1. Supplementary Material.** Supplementary material for: “Synaptic networks shape clinical phenotypes in neurodevelopmental disorders: An integrative clinical, genetic and biological perspective”. Includes Supplementary Figures 1-11 and Supplementary Tables 3-6. *Word file*.

**Supplementary File 2.** Mean brain expression across developmental age for each gene. *Word file*

**Supplementary Table 1.** Summary of the literature review process. *Excel file*.

**Supplementary Table 2**. Mapping of original HPO terms to aggregated categories. *Excel file*.

