## Supplementary File 1 for "Synaptic networks shape clinical phenotypes in neurodevelopmental disorders: An integrative clinical, genetic and biological perspective"

### SUPPLEMENTARY FIGURES

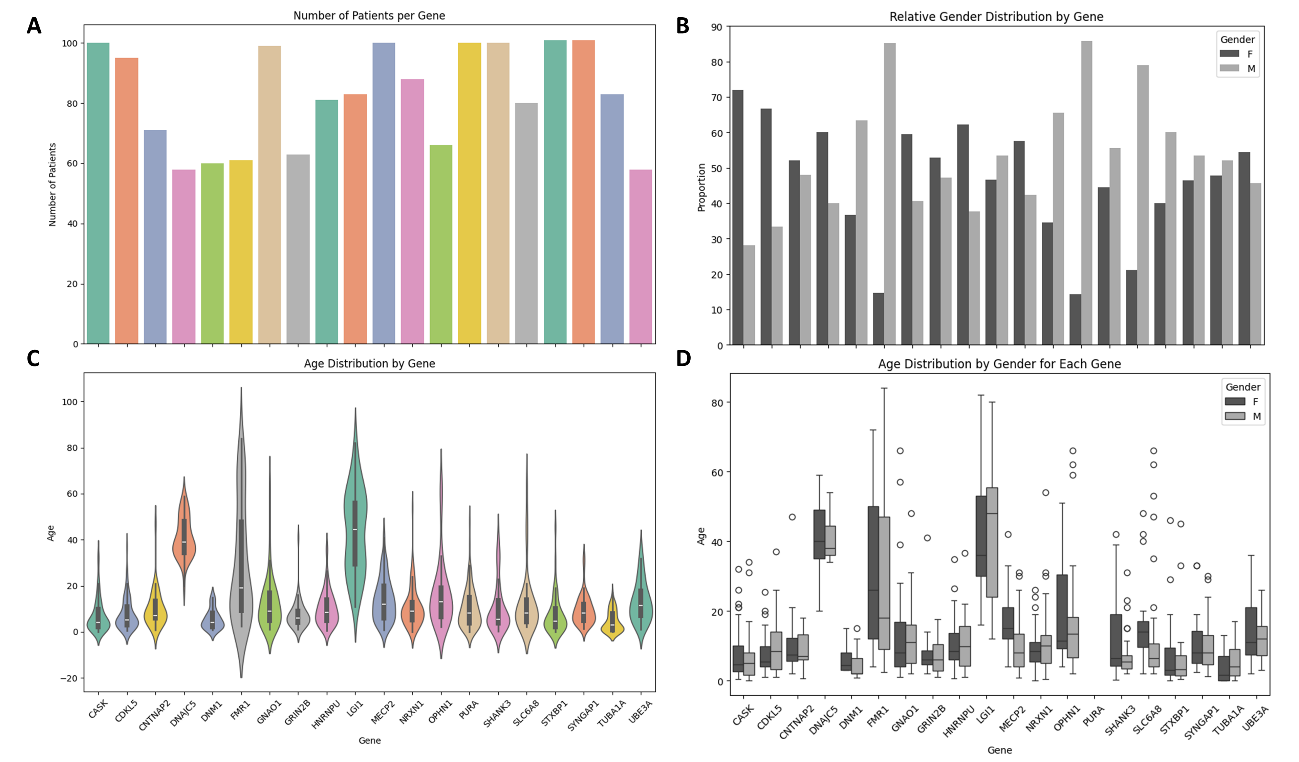

Supplementary Figure 1. Demographic characteristics of the patient cohort, stratified by gene. A) Total number of patients reported per gene. B) Gender distribution (proportion of female and male patients) for each gene. C) Age distribution across all patients per gene. D) Age distribution separated by gender, allowing assessment of sex-specific age patterns.

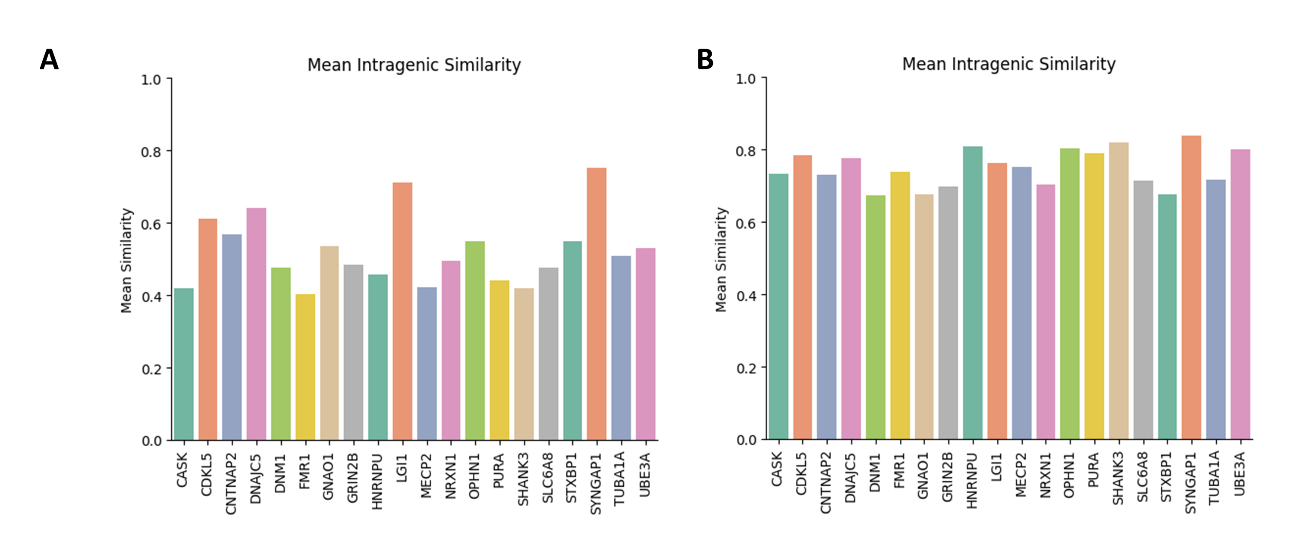

Supplementary Figure 2. Enrichment analysis of neurological symptoms and generalization of non-neurological symptoms across the cohort. A) Intragenic phenotypic similarity scores calculated using raw HPO terms, without any hierarchical grouping, showing variability in symptom profiles across genes. B) Intragenic similarity recalculated after applying an HPO aggregation strategy (see Methods, Section “Symptom Aggregation”), in which individual HPO terms were grouped under broader parent categories. The increase in similarity scores post-aggregation suggests improved phenotypic coherence within genes, especially for those with more heterogeneous clinical descriptions.

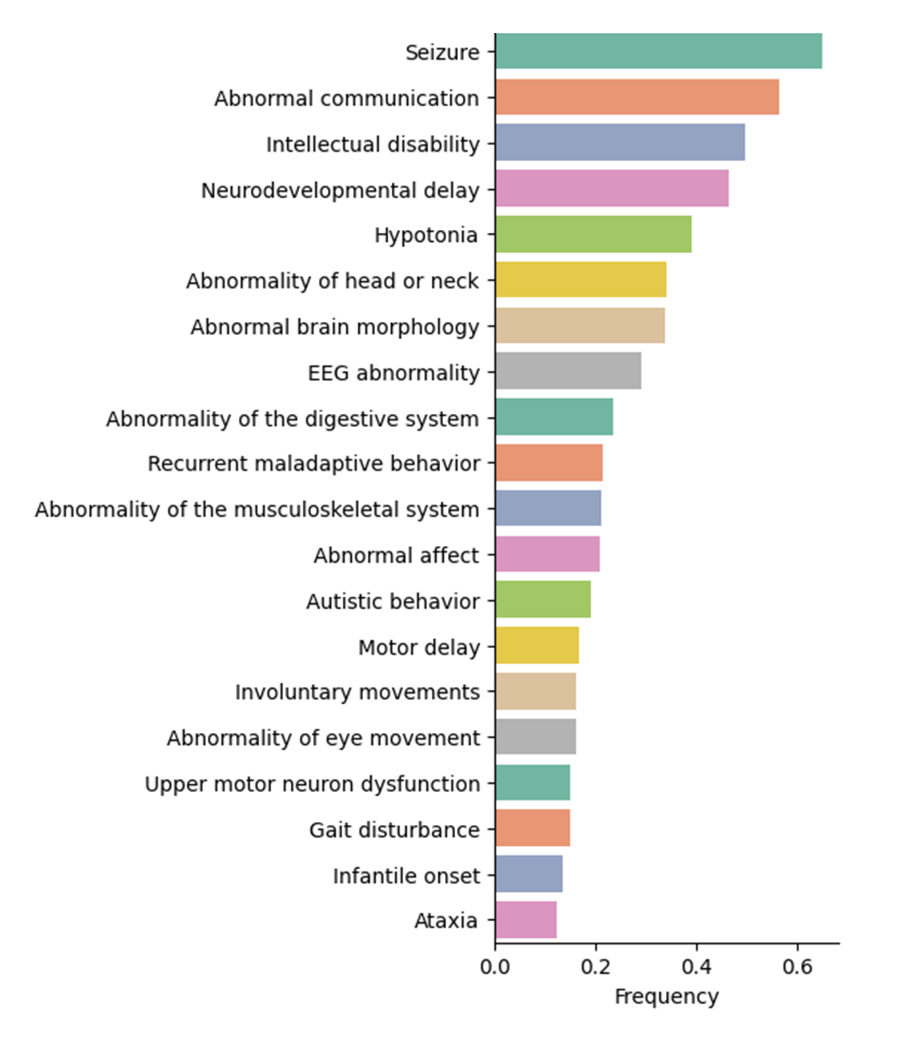

Supplementary Figure 3. Most frequent symptoms across the cohort. Figure shows the 20 most frequently reported symptoms in the full patient cohort, highlighting the predominance of neurological and developmental features. However, among the most frequent features, some non-neurological symptoms were also identified, including abnormalities of the head or neck, musculoskeletal abnormalities, and digestive system disorders.

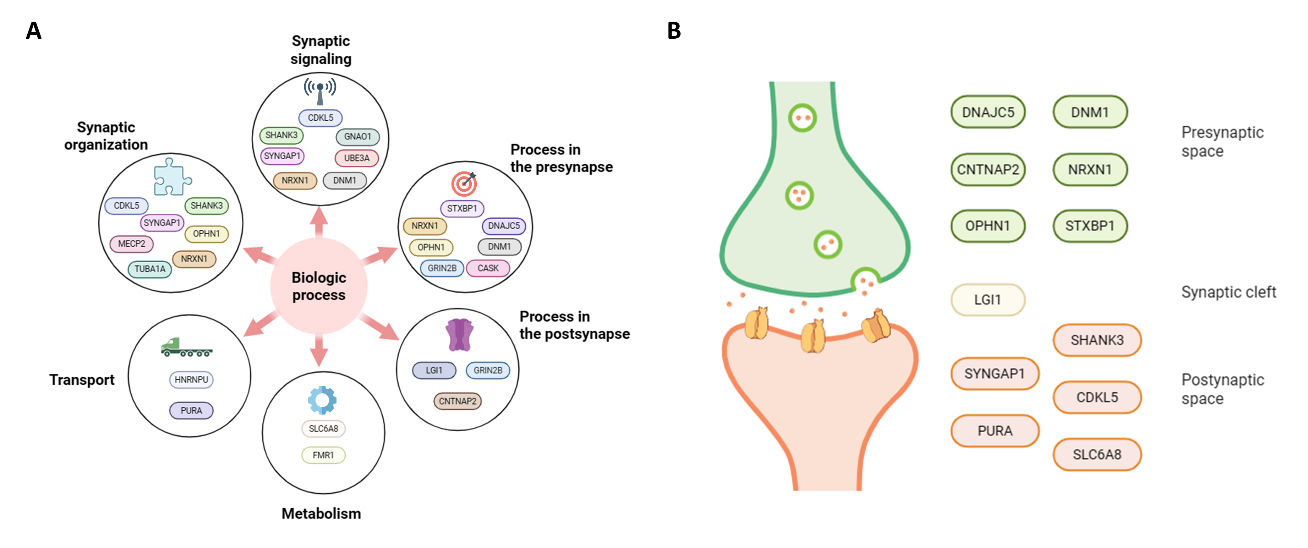

Supplementary Figure 4. Functional classification of the genes selected for the study**. A)** Classification of the genes included in the study based on SynGO-defined biological processes, such as synaptic signaling, synapse organization, metabolism, and transport. **B)** Classification of genes by synaptic localization, distinguishing between presynaptic and postsynaptic compartments.

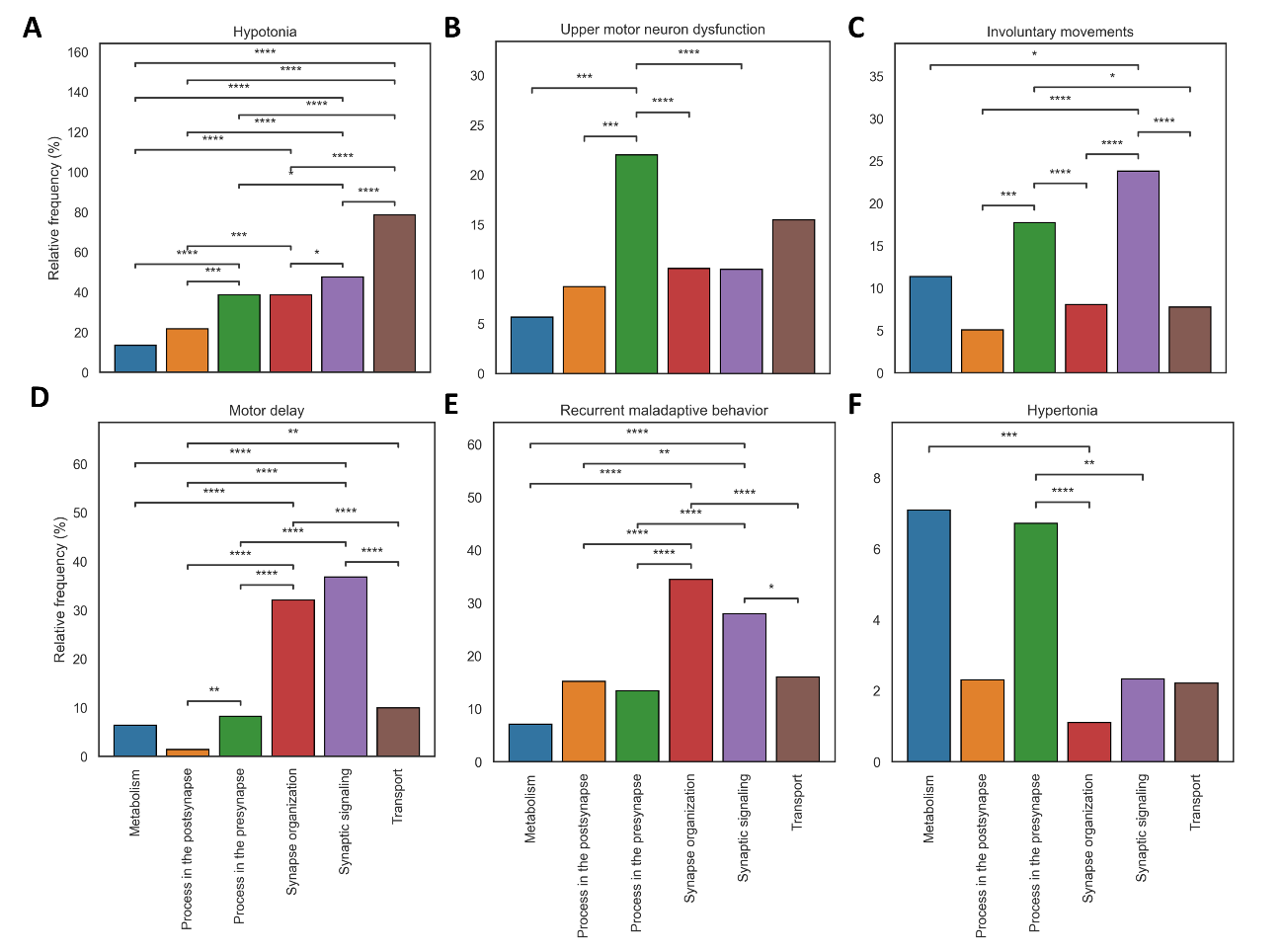

Supplementary Figure 5. Relative frequency (percentage) of symptom presentation across biologic processes (six symptoms selected as examples). **Bars** represent the relative frequency of each symptom for genes within each group. Statistical comparisons were performed using either Fisher’s exact test or **chi-squared test on contingency tables, depending on minimum expected cell counts. Multiple comparisons were corrected using the Bonferroni method.** Horizontal lines indicate pairwise comparisons between groups, with statistical significance denoted by asterisks (p < 0.05: *, p < 0.01: **, p < 0.001: ***, p < 0.0001: ****).

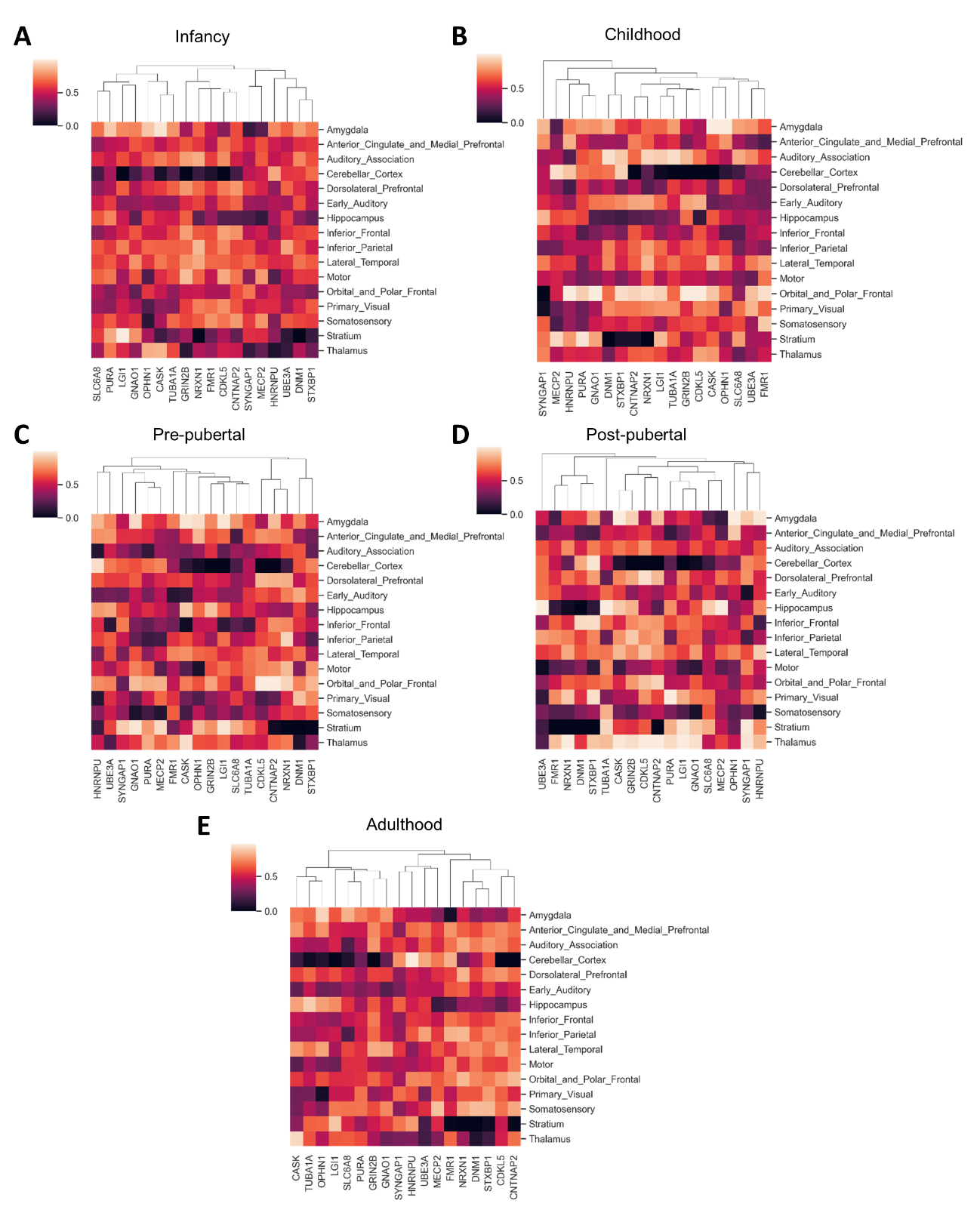

Supplementary Figure 6. Mean expression matrix of genes in the defined developmental stages across brain regions. Brain expression data is obtained from the BrainSpan atlas (see Methods, section “Gene brain expression” for methodological analysis). Developmental groups are A) Infancy (0-2 yo); B) Childhood (3-7 yo); C) Pre-pubertal (8-12.5 yo); D) Post-pubertal (12.5-18 yo); E) Adulthood (>18 yo).

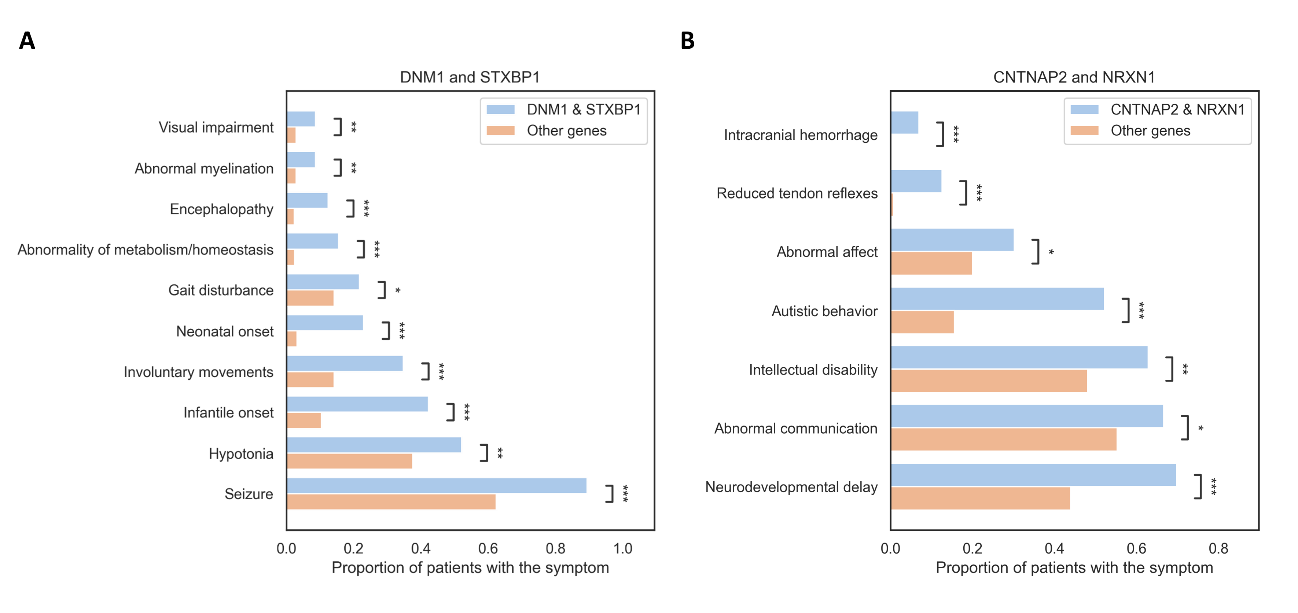

#### Supplementary Figure 7. Proportion of patients presenting significantly more frequent symptoms in the gene pairs.

A) DNM1 and STXBP1, and B) CNTNAP2 and NRXN1, compared to the rest of the genes. Gene pairs with high co-expression across brain development were identified using BrainSpan data. Shown are symptoms with statistically significant differences based on Fisher’s exact test or chi-squared test, with multiple comparisons corrected using the False Discovery Rate (FDR) method. Bars represent the proportion of patients with each symptom in the DNM1 & STXBP1 or CNTNAP2 & NRXN1 group (blue) versus the other genes (orange). Statistical significance is indicated by brackets (adjusted p < 0.05: *, < 0.01: **, < 0.001: ***).

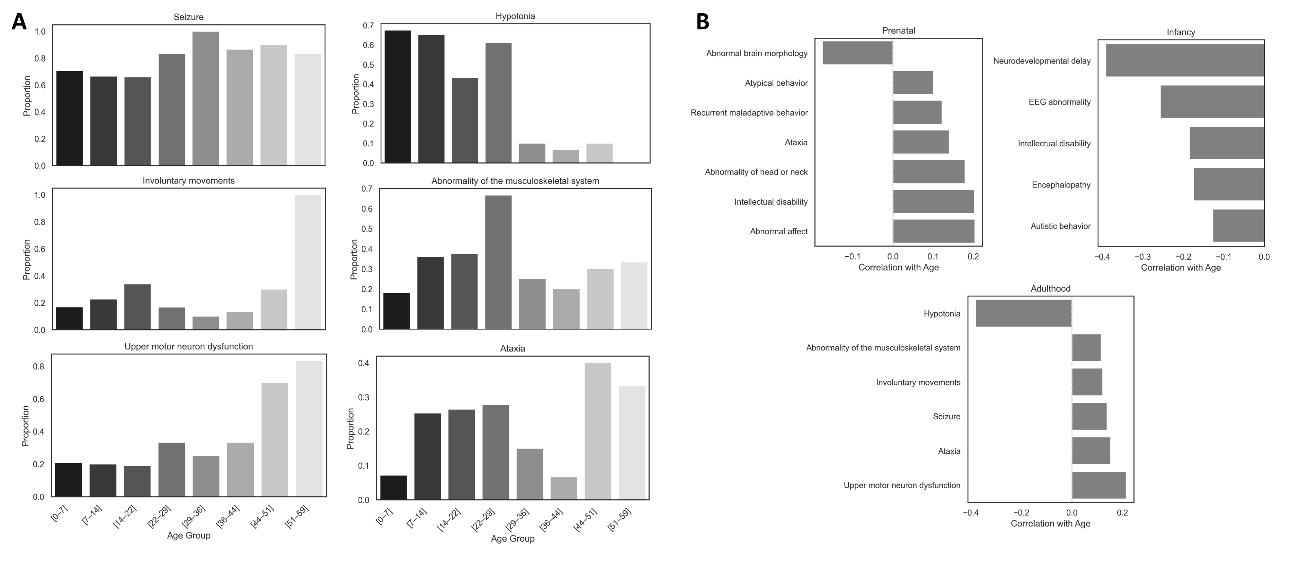

Supplementary Figure 8. Symptom trajectories across age. **A) Proportion of patients with symptoms significantly associated with genes exhibiting peak expression during adulthood and showing a significant correlation with patient age.** Correlations between symptom presence (binary) and age (continuous) were assessed using the point-biserial correlation coefficient. P-values were adjusted for multiple comparisons using the Benjamini-Hochberg false discovery rate (FDR) method, and symptoms with an adjusted p-value (FDR) < 0.05. Each plot illustrates the proportion of patients presenting the symptom across defined age groups: A) Seizure, B) Hypotonia, C) Involuntary movements, D), Abnormality of the musculoskeletal system, E) Upper motor neuron dysfunction, F) Ataxia. B) Correlation between age at symptom onset and symptoms associated with genes of maximum expression in each developmental stage. B) Genes with prenatal peak expression (CASK, SYNGAP1, OPHN1, HNRNPU, TUBA1A, UBE3A, GRIN2B, NRXN1); Genes with infancy peak expression (SLC6A8, FMR1, LGI1, CNTNAP2, CDKL5, GNAO1); Genes with peak expression in post-pubertal/adulthood ages (STXBP1, DNAJC5, PURA, MECP2, DNM1). Only symptoms previously linked to the infant group (based on relative frequency thresholds) were tested. Correlation was computed using the point-biserial correlation coefficient. Multiple comparisons were corrected using the False Discovery Rate (FDR). Only symptoms with adjusted *p* < 0.05 are shown.

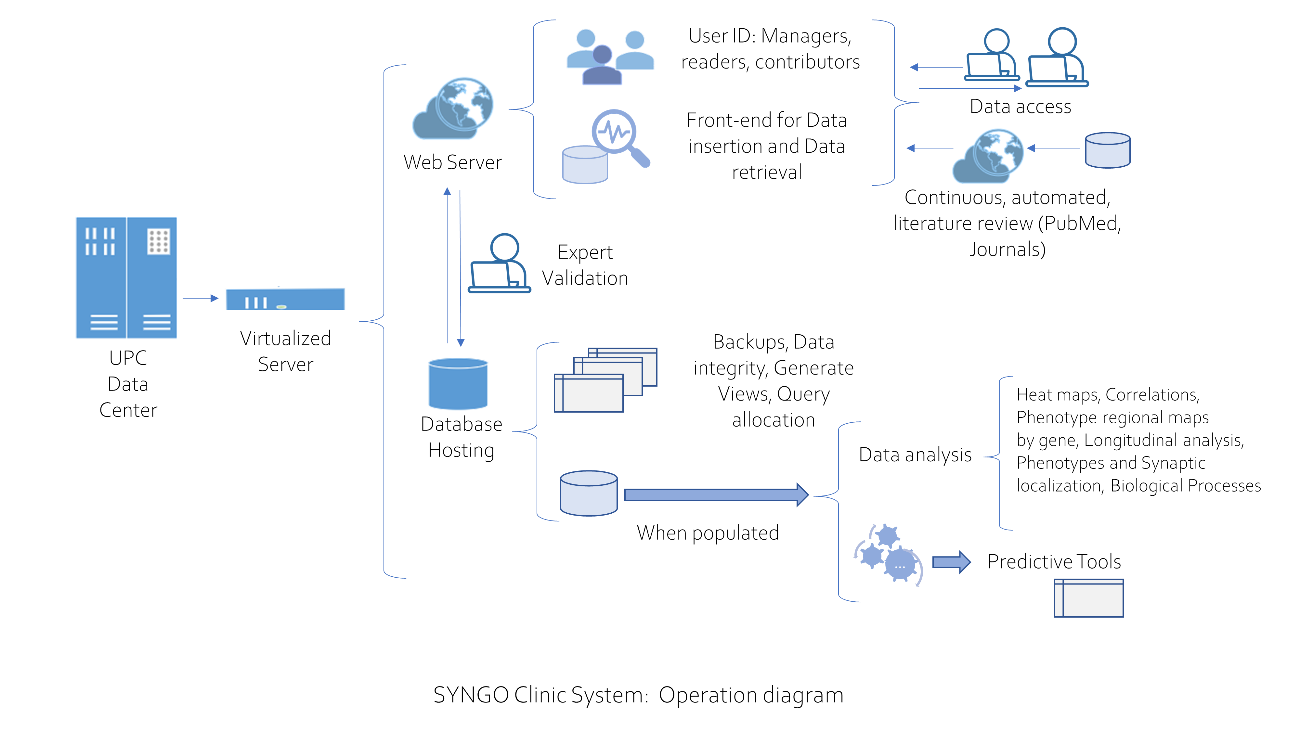

#### Supplementary Figure 9. Conceptual diagram of the clinical-biological integration model proposed “SynGO-Clinic”.

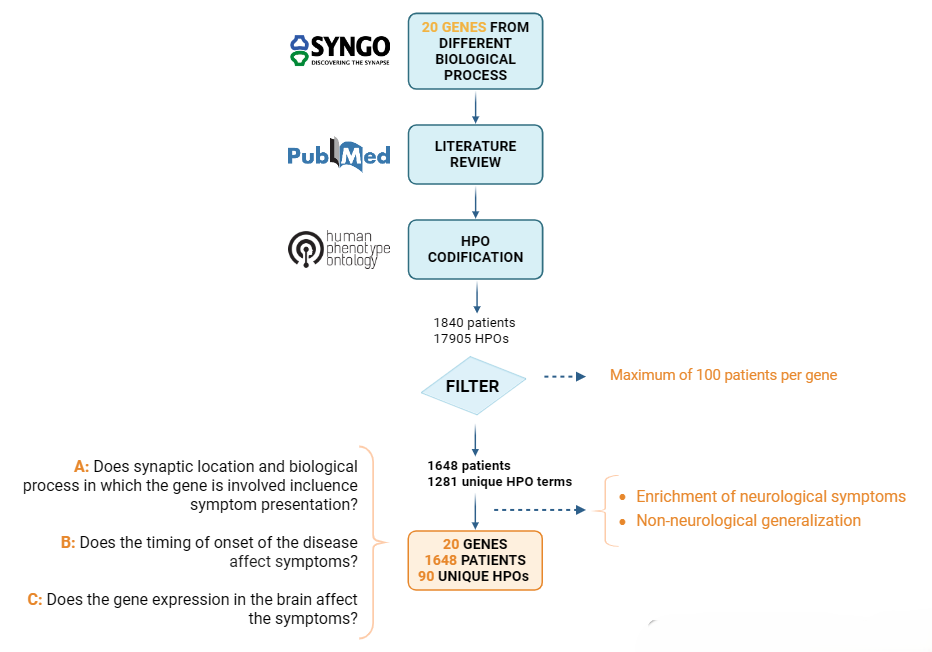

Supplementary Figure 10. Diagram following the data base creation and hypothesis. After gene selection, data is collected by a systemic review and symptoms are codified using HPO terms. A threshold is applied of maximum 100 patients per gene to ensure homogeneity. After that, HPO aggregation into own methods is performed, enabling the enrichment of neurological symptoms. Final database is obtained and the research questions are defined.

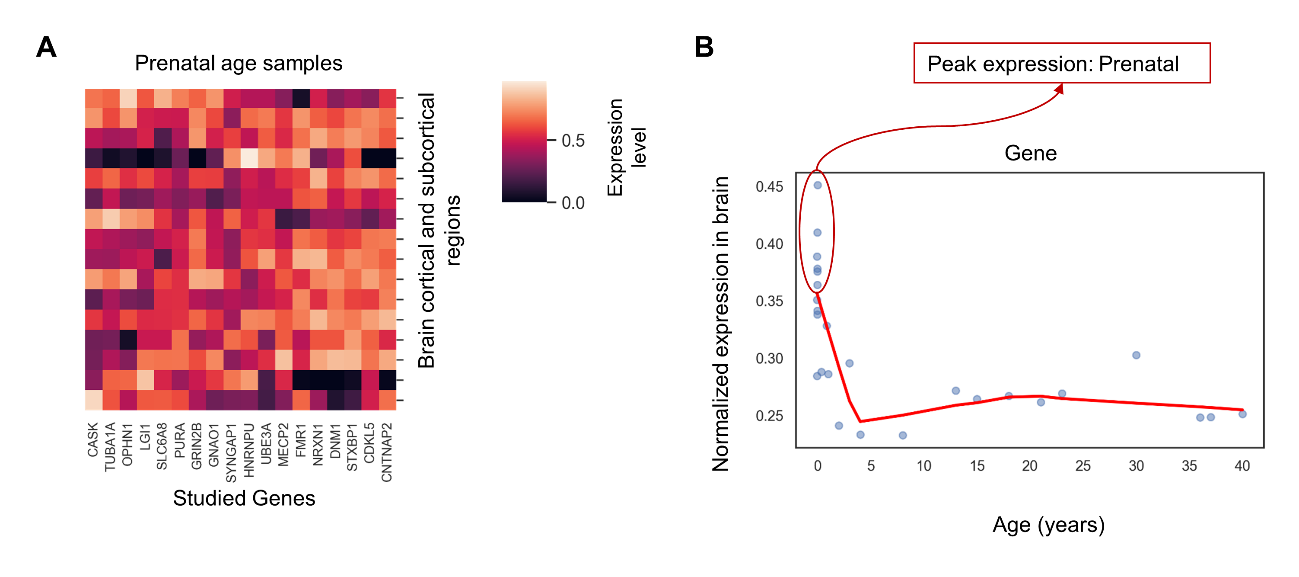

Supplementary Figure 11. Brain expression analysis across development methodology. **A)** For each developmental stage (prenatal, infant, childhood, pre-pubertal, post-pubertal and adulthood), expression matrices are obtained. In these matrices, it is observed how genes are expressed across brain regions in each developmental window. **B**) For each gene, its normalized expression is obtained across development (BrainSpan developmental transcriptome data). The age of peak expression is extracted, and genes are grouped according to their peak expression in the brain.

### SUPPLEMENTARY TABLES

Supplementary Table 1. Summary of the literature review process**.** This table details the systematic review conducted for each gene included in the study. For every gene, the table lists the selected publications, the number of patients reported per article, the number of patients included in our cohort, and the date of search. *Number of patients considered prior to filtering for a minimum number of HPO terms per patient and a maximum of 100 patients per gene ***(Excel file).***

Supplementary Table 2. Mapping of original HPO terms to aggregated categories. This table presents the aggregation strategy applied to Human Phenotype Ontology (HPO) terms used in this study. The left column lists the original HPO terms extracted from clinical reports across the cohort, while the right column indicates the assigned parent term, following the strategy defined in Methods, Section "Symptom Aggregation" ***(Excel file).***

Supplementary Table 3. Correlation between pairs of genes across different developmental stages.

| Developmental age group | GENES | | Correlation (r) | CI95% | p-value |
| --- | --- | --- | --- | --- | --- |
| Infant   (0-2 yo) | CASK | TUBA1A | 0.839582 | [0.59, 0.94] | *p* < 0.001 |
|  | CASK | DNAJC5 | 0.851054 | [0.61, 0.95] | *p* < 0.001 |
|  | DNM1 | STXBP1 | 0.855060 | [0.62, 0.95] | *p* < 0.001 |
|  | CDKL5 | CNTNAP2 | 0.801581 | [0.51, 0.93] | *p* < 0.001 |
|  | TUBA1A | DNAJC5 | 0.879017 | [0.68, 0.96] | *p* < 0.001 |
|  | FMR1 | NRXN1 | 0.832814 | [0.57, 0.94] | *p* < 0.001 |
|  | GRIN2B | CDKL5 | 0.760428 | [0.42, 0.91] | *p* < 0.001 |
|  | DNM1 | NRXN1 | 0.773237 | [0.45, 0.92] | *p* < 0.001 |
|  | CDKL5 | FMR1 | 0.794625 | [0.49, 0.93] | *p* < 0.001 |
|  | CDKL5 | NRXN1 | 0.761899 | [0.43, 0.91] | *p* < 0.001 |
|  | CNTNAP2 | FMR1 | 0.774914 | [0.45, 0.92] | *p* < 0.001 |
|  | CNTNAP2 | NRXN1 | 0.790507 | [0.48, 0.92] | *p* < 0.001 |
| Early childhood  (3-7 yo) | DNM1 | STXBP1 | 0.933736 | [0.82, 0.98] | *p* < 0.001 |
|  | DNM1 | NRXN1 | 0.853948 | [0.62, 0.95] | *p* < 0.001 |
|  | CNTNAP2 | NRXN1 | 0.818065 | [0.54, 0.93] | *p* < 0.001 |
|  | STXBP1 | NRXN1 | 0.847152 | [0.61, 0.95] | *p* < 0.001 |
|  | TUBA1A | OPHN1 | 0.834707 | [0.58, 0.94] | *p* < 0.001 |
|  | CASK | TUBA1A | 0.764425 | [0.43, 0.91] | *p* < 0.001 |
|  | SLC6A8 | DNAJC5 | 0.784174 | [0.47, 0.92] | *p* < 0.001 |
|  | CDKL5 | CNTNAP2 | 0.759052 | [0.42, 0.91] | *p* < 0.001 |
| Pre-pubertal   (8-12.5 yo) | CNTNAP2 | NRXN1 | 0.857684 | [0.63, 0.95] | *p* < 0.001 |
|  | PURA | GNAO1 | 0.769964 | [0.44, 0.92] | *p* < 0.001 |
|  | PURA | MECP2 | 0.754870 | [0.41, 0.91] | 0.75 |
| Post-pubertal   (12.5-18 yo) | CASK | GRIN2B | 0.879547 | [0.68, 0.96] | *p* < 0.001 |
|  | DNM1 | STXBP1 | 0.907549 | [0.75, 0.97] | *p* < 0.001 |
|  | LGI1 | GNAO1 | 0.864646 | [0.65, 0.95] | *p* < 0.001 |
|  | CNTNAP2 | NRXN1 | 0.813942 | [0.53, 0.93] | *p* < 0.001 |
|  | FMR1 | NRXN1 | 0.843158 | [0.6, 0.94] | *p* < 0.001 |
|  | CDKL5 | GNAO1 | 0.793890 | [0.49, 0.93] | *p* < 0.001 |
| Adult  (>18 yo) | DNM1 | STXBP1 | 0.933736 | [0.82, 0.98] | *p* < 0.001 |
|  | DNM1 | NRXN1 | 0.853948 | [0.62, 0.95] | *p* < 0.001 |
|  | CNTNAP2 | NRXN1 | 0.818065 | [0.54, 0.93] | *p* < 0.001 |
|  | STXBP1 | NRXN1 | 0.847152 | [0.61, 0.95] | *p* < 0.001 |
|  | TUBA1A | OPHN1 | 0.834707 | [0.58, 0.94] | *p* < 0.001 |
|  | CASK | TUBA1A | 0.764425 | [0.43, 0.91] | *p* < 0.001 |
|  | SLC6A8 | DNAJC5 | 0.784174 | [0.47, 0.92] | *p* < 0.001 |
|  | CDKL5 | CNTNAP2 | 0.759052 | [0.42, 0.91] | *p* < 0.001 |

Developmental age groups were defined according to the GTEx resource (https://gtexportal.org/home/aboutdGTEx). Gene expression levels were obtained from the BrainSpan Atlas of the Developing Human Brain. For each developmental stage, a mean brain expression matrix was constructed. Pairwise (partial) correlation coefficients were then calculated to assess gene co-expression across developmental windows.

Supplementary Table 4. Mean brain expression across developmental age groups for selected pairs of correlated genes.

|  | Mean expression | | | |
| --- | --- | --- | --- | --- |
| Brain region | **DNM1** | **STXBP1** | **CNTNAP1** | **NRXN1** |
| Amygdala | 0.582927 | 0.543617 | 0.641071 | 0.575721 |
| Anterior Cingulate and Medial Prefrontal | 0.488276 | 0.512666 | 0.650488 | 0.610752 |
| Auditory Association | 0.719994 | 0.682317 | 0.590976 | 0.800400 |
| Cerebellar Cortex | 0.596230 | 0.750686 | 0.051733 | 0.218772 |
| Dorsolateral Prefrontal | 0.554934 | 0.600938 | 0.716593 | 0.705592 |
| Early Auditory | 0.580509 | 0.567343 | 0.581262 | 0.648317 |
| Hippocampus | 0.358474 | 0.279930 | 0.319386 | 0.301956 |
| Inferior Frontal | 0.602799 | 0.595646 | 0.731530 | 0.631217 |
| Inferior Parietal | 0.621325 | 0.593263 | 0.662986 | 0.839782 |
| Lateral Temporal | 0.721226 | 0.625244 | 0.664443 | 0.766355 |
| Motor | 0.543644 | 0.515902 | 0.655048 | 0.586522 |
| Orbital and Polar Frontal | 0.618493 | 0.631463 | 0.820286 | 0.746131 |
| Primary Visual | 0.643667 | 0.757493 | 0.556176 | 0.690387 |
| Somatosensory | 0.651115 | 0.657067 | 0.692209 | 0.632058 |
| Stratium | 0.001190 | 0.054230 | 0.066558 | 0.001636 |
| Thalamus | 0.226554 | 0.319674 | 0.657336 | 0.523735 |
| Correlation | **0.933, *p* < 0.001** | | **0.878, *p* < 0.001** | |

Expression values were obtained from the BrainSpan Atlas and normalized across samples and genes. The Pearson correlation of mean expression across regions was r = 0.933, p = 1.27 × 10⁻⁷ for *DNM1* and *STXBP1*, and r = 0.870, p = 7.75 × 10⁻⁶ for *CNTNAP2* and *NRXN1*, indicating strong co-expression across brain development.

Supplementary Table 5. Symptoms associated with each gene group categorized by the developmental stage of peak brain expression.

| Group of gene maximum expression | Symptom | Frequency (%) | p-value |
| --- | --- | --- | --- |
| Prenatal | Abnormality of the eye | 17 | *p* < 0.001 |
|  | Abnormal affect | 26.56 | *p* < 0.001 |
|  | Abnormality of the musculoskeletal system | 25.31 | *p* < 0.001 |
|  | Abnormal brain morphology | 52.03 | *p* < 0.001 |
|  | Upper motor neuron dysfunction | 15.16 | *p* < 0.001 |
|  | Abnormality of head or neck | 47.34 | *p* < 0.001 |
|  | Hypotonia | 37.18 | *p* < 0.001 |
|  | Abnormality of limbs | 13.44 | *p* < 0.001 |
|  | Ataxia | 15 | *p* < 0.001 |
|  | Abnormality of eye movement | 24 | *p* < 0.001 |
|  | Atypical behavior | 15.78 | *p* < 0.001 |
|  | Autistic behavior | 25 | *p* < 0.001 |
|  | Developmental stagnation/regression | 8.4375 | *p* < 0.001 |
|  | Intellectual disability | 65 | *p* < 0.001 |
|  | Recurrent maladaptive behavior | 20.94 | *p* < 0.001 |
|  | Neurodevelopmental delay | 66.25 | *p* < 0.001 |
|  | Motor delay | 21.41 | *p* < 0.001 |
|  | Sleep abnormality | 13.75 | *p* < 0.001 |
| Infant (1-3 years) | Abnormal affect | 18.81 | *p* < 0.001 |
|  | Autistic behavior | 16.77 | *p* < 0.001 |
|  | Encephalopathy | 5.52 | 0.005 |
|  | Seizure | 71.16 | *p* < 0.001 |
|  | Intellectual disability | 40.9 | *p* < 0.001 |
|  | EEG abnormality | 37.63 | *p* < 0.001 |
|  | Neurodevelopmental delay | 42.33 | *p* < 0.001 |
| Adult (>18 years) | Abnormality of the musculoskeletal system | 24.82 | *p* < 0.001 |
|  | Upper motor neuron dysfunction | 26.73 | *p* < 0.001 |
|  | Involuntary movements | 21.72 | *p* < 0.001 |
|  | Hypotonia | 51.55 | *p* < 0.001 |
|  | Abnormality of limbs | 8.83 | *p* < 0.001 |
|  | Ataxia | 18.14 | *p* < 0.001 |
|  | Seizure | 74.94 | *p* < 0.001 |
|  | Gait disturbance | 27.45 | *p* < 0.001 |
|  | EEG abnormality | 38.42 | *p* < 0.001 |

Genes were grouped into prenatal, infant (1–3 years), and adult (>18 years) categories based on the age at which their maximum expression was observed in the BrainSpan Atlas. The table lists the symptoms most frequently associated with each group, reflecting characteristic clinical profiles linked to the timing of gene brain expression during development. Symptom frequency is reported as a percentage, and *p*-values correspond to comparisons between groups. Statistical differences were assessed using Fisher’s exact test or chi-squared test, depending on expected cell counts. Only symptoms meeting the threshold for significance (*p-adjusted* < 0.05) and passing group-level dominance criteria (see Methods, Section “Symptom Frequency Analysis”) are included.

Note: Genes with prenatal peak expression (CASK, SYNGAP1, OPHN1, HNRNPU, TUBA1A, UBE3A, GRIN2B, NRXN1); Genes with infancy peak expression (SLC6A8, FMR1, LGI1, CNTNAP2, CDKL5, GNAO1); Genes with peak expression in post-pubertal/adulthood ages (STXBP1, DNAJC5, PURA, MECP2, DNM1).

##### Supplementary Table 6. Criteria for gene dominance.

| Group size | Dominance criteria |
| --- | --- |
| 2–3 genes | If any single gene ≥ 70%, it is considered gen x 10-dominance |
| > 3 genes | If the sum of the top 2 genes ≥ 80%, it is considered gen x 10-dominance |
