## Supplementary figures and images for "Synaptic networks shape clinical phenotypes in neurodevelopmental disorders: An integrative clinical, genetic and biological perspective"

### Supplementary File 2

**Supplementary File 2. Mean brain expression across developmental age for each gene.**

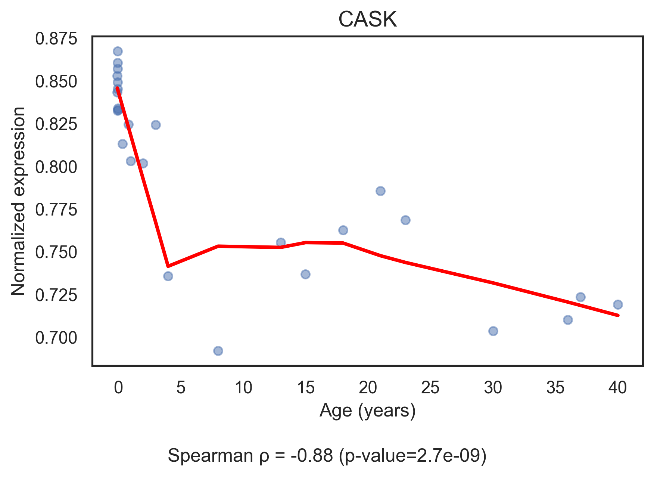

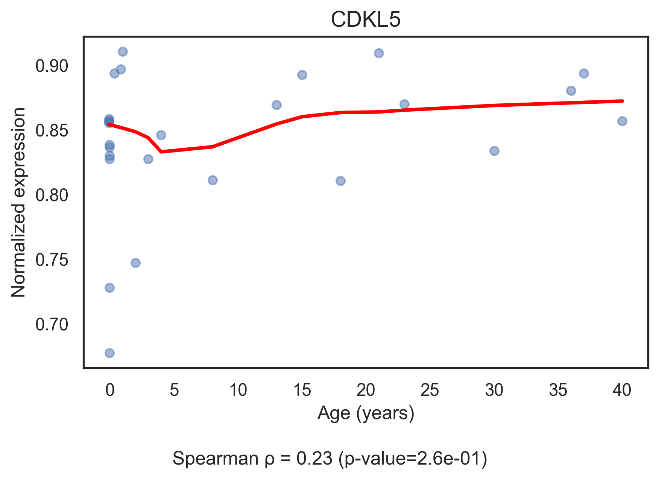

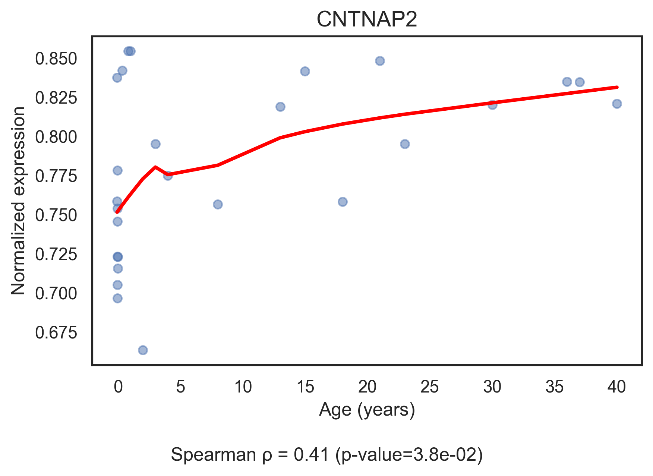

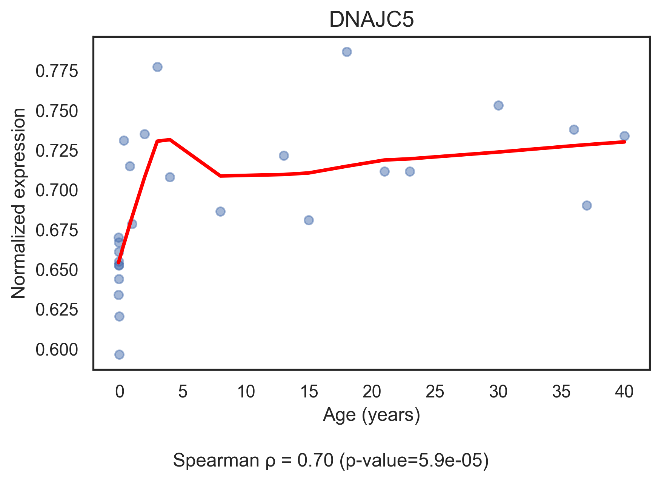

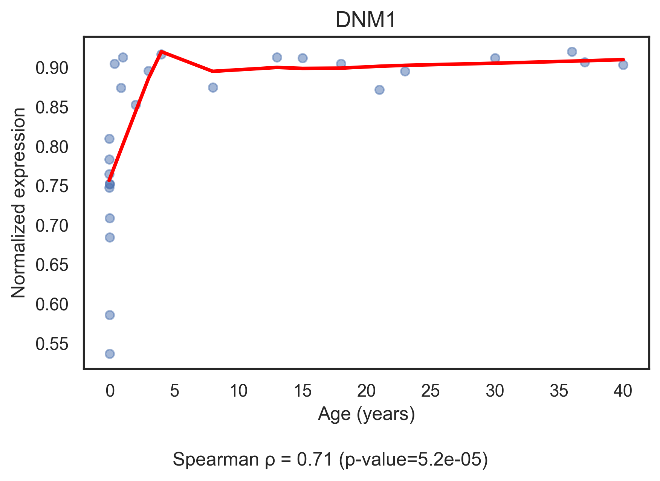

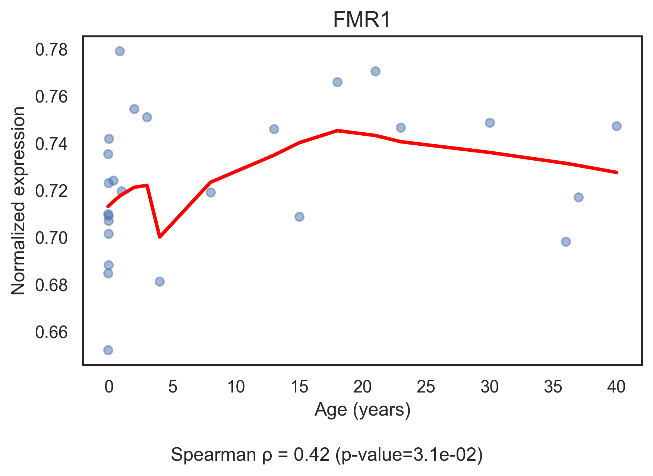

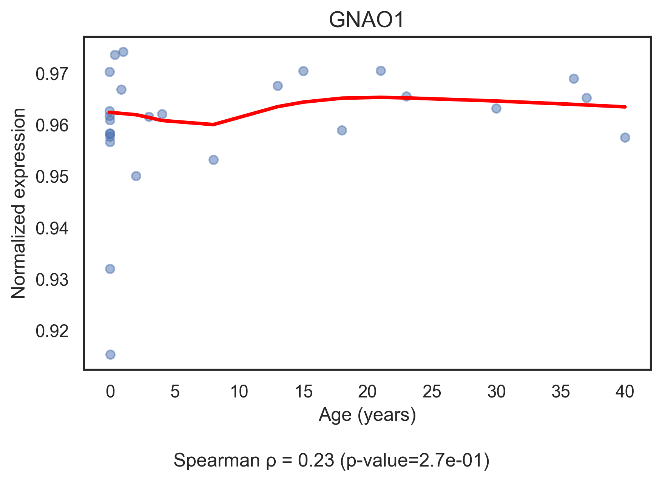

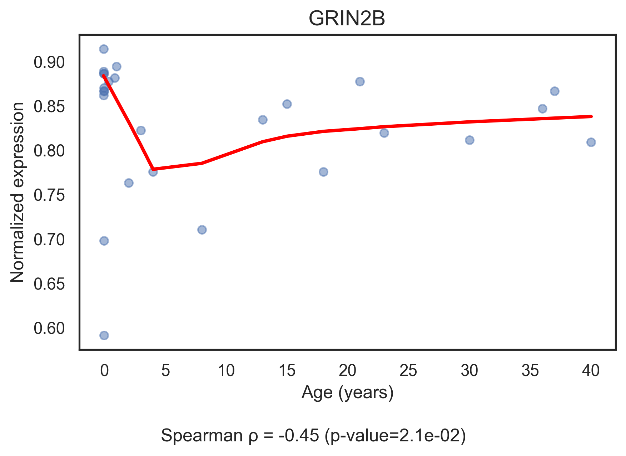

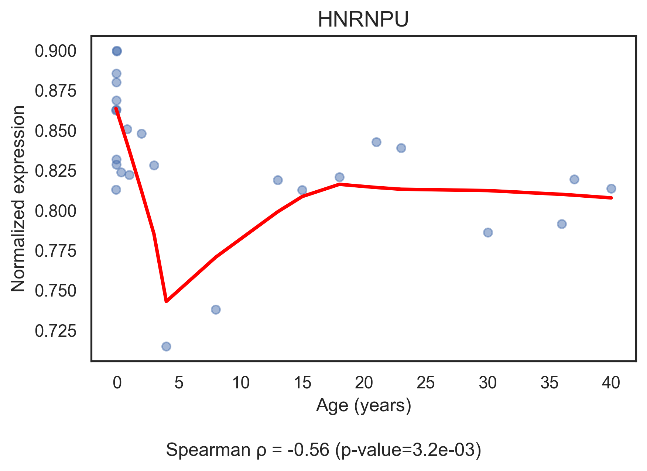

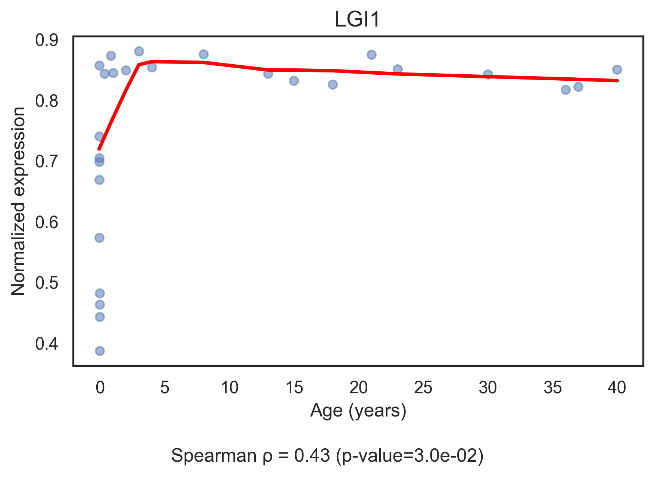

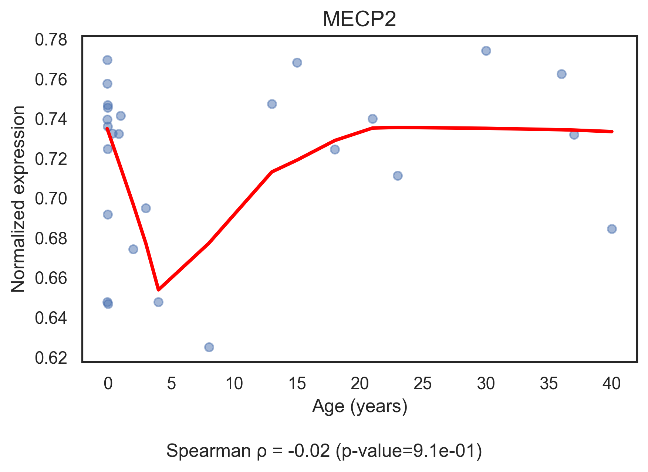

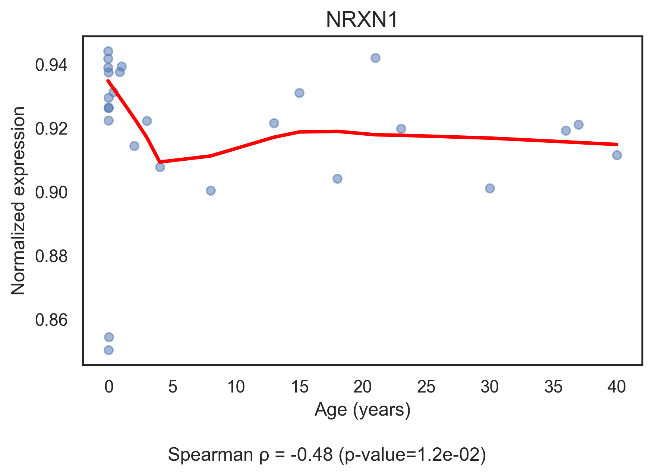

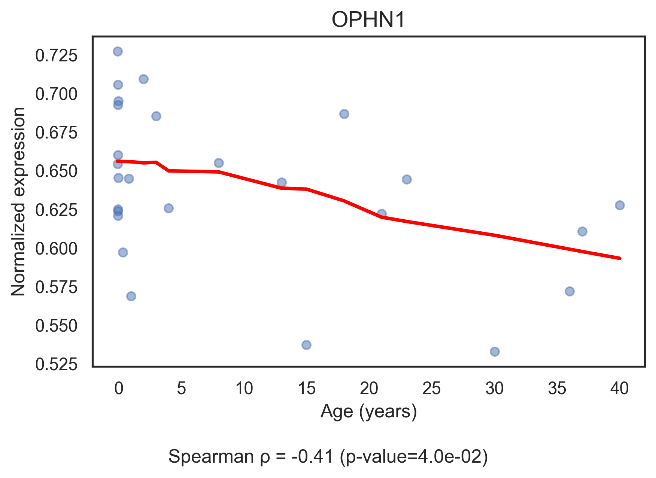

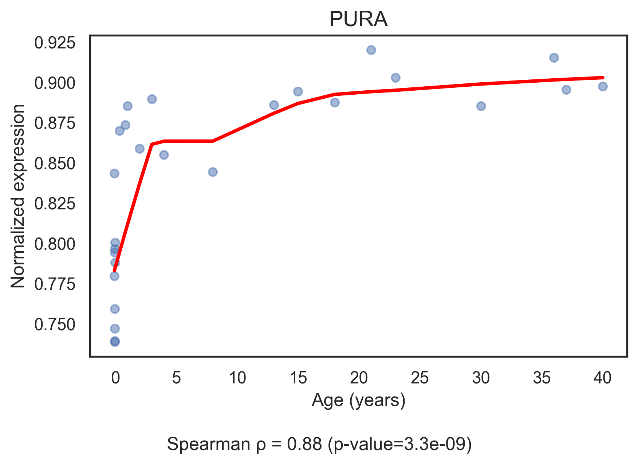

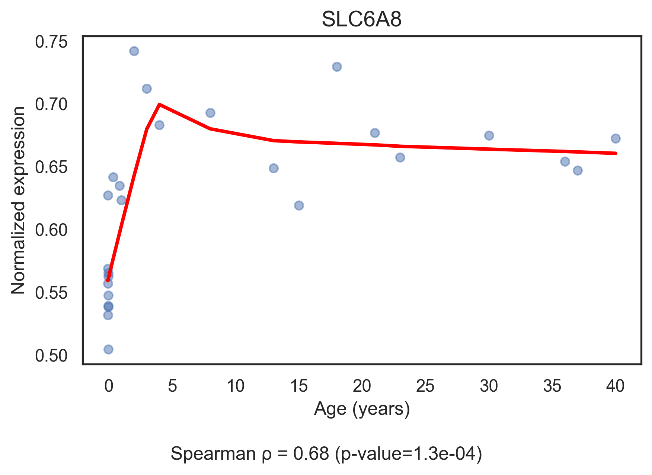

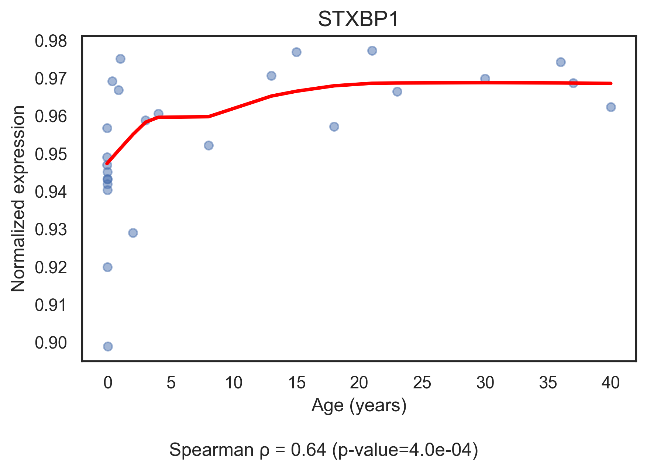

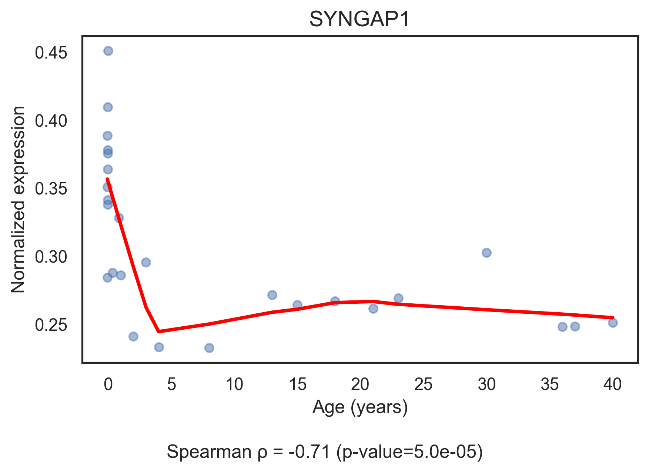

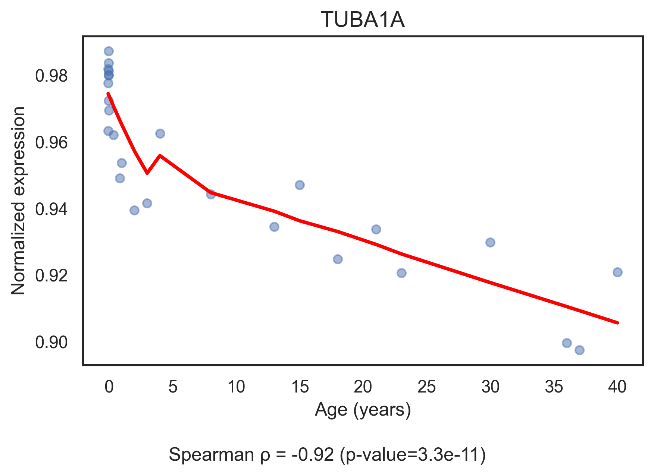

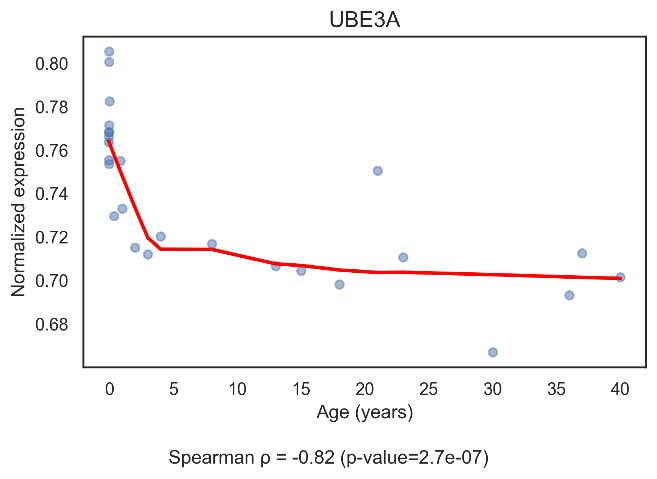
